# Loss of *Kmt2c*/*d* promotes gastric cancer initiation and confers vulnerability to mTORC1 inhibition and anti-PD1 immunotherapy

**DOI:** 10.1101/2025.03.27.645747

**Authors:** Naitao Wang, Dan Li, Tao Zhang, Mohini R. Pachai, Woo Hyun Cho, Makhzuna N. Khudoynazarova, Dana M. Schoeps, Yudi Bao, Marion Liu, Laura Tang, Janjigian Yelena, Ping Chi, Yu Chen

**Author notes:** These authors contributed equally to this work (Naitao Wang, Dan Li, Tao Zhang).

## Abstract

*KMT2C* and *KMT2D* (*KMT2C/D*) are frequently mutated in gastric adenocarcinoma, yet their function in cancer initiation remains poorly understood. In this study, based on the observation that loss-of-function mutations of *KMT2C* and *KMT2D* are enriched and co-occur in gastric adenocarcinoma, we developed genetically engineered mouse models to selectively knock out *Kmt2c* and *Kmt2d* in gastric epithelial cells with *Tmprss2-CreER^T2^*. Through histological staining and single-cell RNA sequencing, we observed that *Kmt2c/d* loss led to nuclear dysplasia and expansion of cells with mixed gastric lineage markers. When combined with Pten deletion, Kmt2c/d loss drove rapid development of muscle-invasive gastric adenocarcinoma as early as 3 weeks post Cre-mediated gene deletion. The adenocarcinoma exhibited decreased expression of gastric lineage markers and increased expression of intestinal differentiation markers, phenocopying human gastric adenocarcinoma. *Kmt2c/d* knockout reduced protein synthesis but upregulated transcription of ribosomal proteins, rendering sensitivity to mTORC1 inhibitors. Additionally, *Kmt2c/d* knockout increased MHC-I molecule expression and enhanced antigen presentation. Combination of mTROC1 inhibition and anti-PD1 immunotherapy significantly suppressed tumor growth in immune-competent mice. Together, these findings reveal the role of *Kmt2c*/*d* loss in gastric cancer initiation and suggest the potential therapeutic strategies for *KMT2C/D*-deficient gastric cancer.

## Introduction

Gastric or stomach adenocarcinoma (STAD) is the fifth most common cancer and the fourth most common cause of cancer death worldwide [1]. Gastric cancer exhibits histological and molecular heterogeneity, correlating with diverse genetic mutations and clinical outcomes [2]. Based on molecular profiles, STAD can be classified into four major subtypes: microsatellite instability (MSI), Epstein–Barr virus-positive (EBV), genomically stable (GS), and chromatin instability (CIN) [3, 4]. Histologically, MSI, EBV, and CIN tumors exhibit intestinal type and GS tumors exhibit diffuse type by Lauren classification [5]. *PIK3CA* hotspot mutations are the most prevalent in EBV subtype, loss of *CDH1* is the dominant mutation in GS subtype, and *TP53* loss-of-function (LOF) mutations are the leading alteration in CIN subtype. While dysfunction of the mismatch repair (MMR) system is the primary cause of hypermutagenesis in MSI subgroup, the drivers of cancer initiation in this subtype remain largely unexplored. In the MSI subgroup, mutations in the RAS and phosphoinositide 3-kinase (PI3K) pathway, and epigenetic modifiers, such as *KMT2D*, *ARID1A*, and *KMT2C*, represent top genetic alterations [3, 4]. Recent studies using human stomach organoid and genetically engineered mouse models (GEMMs) have demonstrated that loss of *ARID1A* promotes cancer progression and impairs tumor differentiation [6, 7]. However, the roles of *KMT2C* and *KMT2D* alterations in STAD remain unclear.

KMT2C and KMT2D are members of type 2 histone lysine methyltransferase (KMT2) family [8, 9]. The primary catalytic function of KMT2C/D is to mediate mono- and di-methylation of histone 3 lysine 4 (H3K4me1 and H3K4me2) at active enhancers and a subset of CpG-low promoters [10, 11]. Previous studies have demonstrated the tumor-promoting effects of *KMT2C/D* LOF mutations in lymphomas, urothelial cancer, lung cancer, and breast cancer [10, 12, 13, 14, 15, 16]. In STAD, most *KMT2C/D* mutations are LOF mutations, such as frameshift and nonsense mutations that lead to protein truncations [17], suggesting that the subsequent reduced H3K4 methylation and dysregulated gene expression may contribute to tumorigenesis in gastric cancer as well. Consistently, inhibiting lysine-specific demethylase 1 (LSD1), one of the demethylases of H3K4 methylations, has been shown to suppress gastric cancer cell growth in cell culture and xenograft models [18]. Since LSD1 inhibition increases H3K4 methylation, we hypothesize that loss of *KMT2C/D* and the consequent decrease of H3K4 methylation may promote cancer cell growth [18]. To date, only a few studies have explored the molecular features and clinical implications of *KMT2C/D* mutations in the context of immunotherapy [19].

To characterize the functional consequences of *KMT2C/D* loss in gastric cancer, we developed genetically engineered mouse models (GEMMs) to conditionally knock out *Kmt2c*/*d* in gastric epithelial cells using *Tmprss2-CreER^T2^*. We investigated the histological and transcriptional changes and explored the underlying mechanisms and therapeutic opportunities.

## Results

### Co-occurrence of *KMT2C* and *KMT2D* LOF mutations and PI3K pathway alterations in STAD

To examine the mutational landscape of *KMT2C* and *KMT2D*, we analyzed The Cancer Genome Atlas (TCGA) dataset, focusing on microsatellite instability (MSI)-enriched cancer types, e.g., stomach, colorectal and endometrial adenocarcinoma [3]. In STAD, *KMT2C* mutations were identified in 17.4% of samples, and *KMT2D* mutations were found in 23.2% of samples, with a substantial enrichment in the MSI subgroup (**Figure 1A, B**). Most *KMT2C* and *KMT2D* mutations were bona fide LOF (i.e., truncation and splice site mutations) instead of missense or inframe mutations of unknown significance, suggesting that *KMT2C/D* LOF mutations may be positively selected in cancer progression (**Figure 1C**). Given the high mutational burden of the MSI subgroup due to dysfunction of the MMR system, we analyzed the mutational landscape of other epigenetic modifiers and several comparably large genes. In STAD, we observed that high percentage of *ARID1A* (82.1%), *KMT2D* (69.1%), and *KMT2C* (54.9%) mutations were LOF (**Figure 1C**). In contrast, the majority of mutations of other epigenetic modifiers were missense and the LOF mutation rate of DMD, the largest known human gene, was only 14.3% (**Figure 1C**).

**Figure 1.**
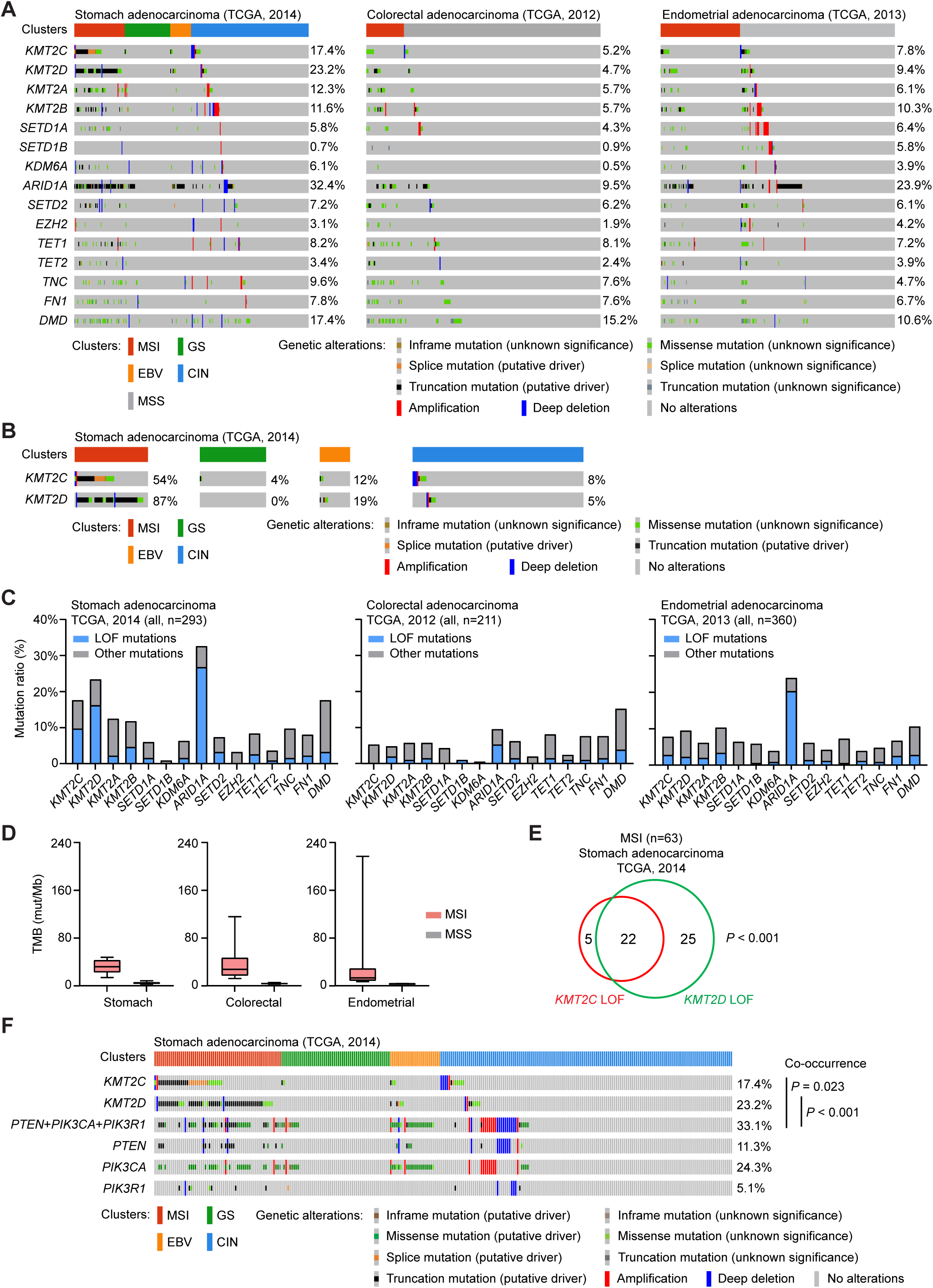
Co-occurrence of *KMT2C/D* LOF mutations and PI3K pathway alterations in STAD. A, OncoPrint of selected chromatin modifying genes and several other large genes in TCGA datasets of stomach adenocarcinoma (2014), colorectal adenocarcinoma (2012), and endometrial adenocarcinoma (2013). Samples in stomach adenocarcinoma were sorted as microsatellite instability (MSI), genomically stable (GS), Epstein-Barr virus-positive (EBV), and chromatin instability (CIN) groups. Samples in colorectal adenocarcinoma and endometrial adenocarcinoma were sorted as MSI and microsatellite stable (MSS). B, OncoPrint of *KMT2C* and *KMT2D* in the TCGA stomach adenocarcinoma dataset. C, Percent of loss-of-function (LOF) mutations and other mutations in stomach adenocarcinoma, colorectal adenocarcinoma, and endometrial adenocarcinoma. Splice and nonsense mutations that lead to protein truncations were considered as LOF mutations. D, Tumor mutational burden in hypermutated and non-hypermutated cancer samples. The center line represents the median, the box limits represent the upper and lower quartiles, and the minimum and maximum whiskers represent the 10^th^ and 90^th^ percentiles, respectively. E, Venn diagram showing the overlap of *KMT2C* and *KMT2D* LOF mutations in MSI STAD samples. F. OncoPrint of *KMT2C*, *KMT2D*, and mutations of PI3K signaling (including *PTEN*, *PIK3CA*, and *PIK3R1*). Mutual co-occurrences were observed between pooled mutations of PI3K signaling and *KMT2C* or *KMT2D*.

To determine whether the positive selection of *KMT2C/D* LOF mutations is specific to STAD, we analyzed their mutational profiles in colorectal and endometrial adenocarcinoma, two other cancer types with a high proportion of MSI tumors [20, 21]. Notably, we observed positive enrichment of LOF mutations in *ARID1A* but not in *KMT2C* or *KMT2D* in these two cancer types (**Figure 1C**). The mutational rates of other genes, such as *TNC*, *FN1*, and *DMD*, were comparable across these cancer types, and the mutational burdens in MSI samples were similar (**Figure 1C, D**). Additionally, we observed the mutual co-occurrence of *KMT2C* and *KMT2D* LOF mutations in MSI subgroup of STAD (**Figure 1E**). These data suggest that *KMT2C* and *KMT2D* LOF mutations may cooperatively contribute to tumorigenesis in STAD.

Dysregulation of the PI3K pathway (e.g., LOF mutations in *PTEN* and *PIK3R1*, and gain-of-function mutations in *PIK3CA*) is common in gastric cancers (**Figure 1F**), particularly in the MSI and EBV subgroups in both TCGA and The Asian Cancer Research Group (ACRG) cohorts [3, 4]. Moreover, *PTEN* inactivation has been shown to accelerate gastric cancer progression [22, 23]. In the TCGA dataset, we observed a high frequency of PI3K pathway alterations, most commonly through *PTEN* LOH mutations and *PIK3CA* hotspot gain-of-function mutations in the MSI subgroup (**Figure 1F**). Additionally, there was a significant co-occurrence of *KMT2C/D* mutations with PI3K pathway alterations (**Figure 1F**), indicating a possible synergistic role in gastric cancer pathogenesis.

### *Kmt2c/d* knockout cooperates with *Pten* loss to induce muscle-invasive gastric cancer

To investigate the functional consequences of *Kmt2c*, *Kmt2d* and *Pten* loss, we utilized *Tmprss2-CreER^T2^-IRES-nlsEGFP* (referred to as *Tmprss2-CreER^T2^* hereafter) to mediate tamoxifen-induced deletion of floxed *Kmt2c*, *Kmt2d,* and *Pten* alleles in epithelial cells of prostate, bladder, and gastrointestinal tract [10, 24, 25]. Consistent with other stomach epithelium specific Cre [26], *Tmprss2-CreER^T2^* induced LoxP recombination in the majority epithelial cells but no stromal cells in *Rosa26-CAG-LSL-EYFP* (TY) mice (**Figure 2A, B**).

**Figure 2.**
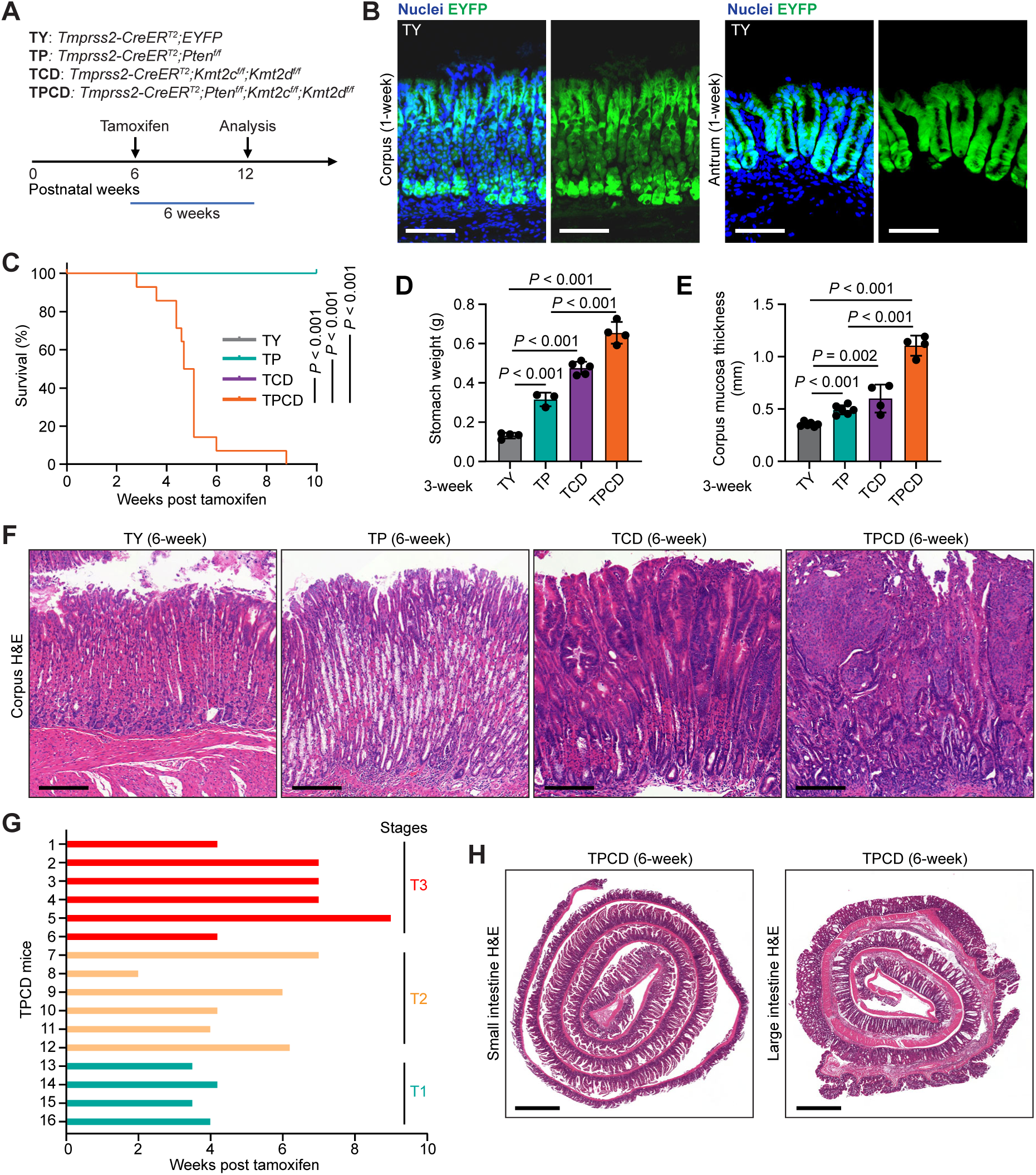
*Kmt2c/d* knockout cooperates with *Pten* loss to induce muscle-invasive stomach cancer. A, Schematic of mouse models: two doses of tamoxifen (3 mg × 2) were injected intraperitoneally with a 48-hour interval. B, Representative immunofluorescence (IF) staining of EYFP in stomach tissues from *Tmprss2-CreER^T2^;Rosa26-LSL-EYFP mice*. Nuclei were counterstained with DAPI. Scale bar, 200 µm. C, Kaplan-Meier plots showing the survival of mice after gene knockout. D, Stomach weight of the indicated genetically engineered mouse model (GEMM) six weeks after tamoxifen injection. Data are presented as mean ± SD and analyzed with two-tailed t-test. E, Thickness in corpus mucosa measured by microscopy of hematoxylin and eosin (H&E) staining section. Each dot represents the averaged thickness of 5 random fields from one mouse. Data are presented as mean ± SD and analyzed with two-tailed t-test. F, Representative H&E staining of stomach tissues after tamoxifen administration. Scale bar, 200 µm. G, Pathological staging of stomach cancer progression based on infiltration of tumor cells in TPCD mice. H, Representative H&E staining of small and large intestines in TPCD mice. Scale bar, 1 mm.

We then crossed the *Tmprss2-CreER^T2^* with *Kmt2c^f/f^* (TC)*, Kmt2d^f/f^* (TD), the combination (TCD), and all with *Pten^f/f^*to generate TP, TPC, TPD and TPCD mouse models, respectively. The deletion efficiency of the conditional *Kmt2c* and *Kmt2d* alleles in each model was validated by *in situ* hybridization (BaseScope™) of the floxed exons and immunohistochemistry (IHC) against H3K4me1 (**Figure S1A**). To assess the deletion efficiency of the conditional *Pten* allele, we performed IHC against PTEN and serine 473 phosphorylated AKT (p-AKT) and observed widespread epithelial PTEN loss in the stomachs of TP and TPCD mice (**Figure S1A**).

Our prior work showed that TC, TD and TCD mice had normal lifespans, whereas TPC and TPD male mice lived to ∼6 and ∼12 months, respectively, due to urothelial cancer leading to urinary obstruction and renal failure. In contrast, TPCD mice survived for only 6 weeks after tamoxifen administration due to stomach cancer and malnutrition [10]. Here, we confirmed the post-tamoxifen survival rates of TY, TP, TCD, and TPCD mice in an independent cohort (**Figure 2C**) and collected stomachs for histology analysis at several time points in concordance with their survival (**Table S1**).

We examined the stomach tissues at the 12-month timepoint for TC and TD groups and at the 6-month timepoint for TPC and TPD groups (**Figure S2A**). Only minor histological changes were observed in the gastric mucosa of TC and TD groups at 12 months (**Figure S2B**), whereas both TPC and TPD groups exhibited dysplasia at 6 months in TPD mice. In TPD mice, there were focal lesions that progressed to carcinoma *in situ* (CIS) in TPD mice (**Figure S2C, D**). In TPD mice, IHC showed that regions of dysplasia and CIS exhibited loss of PTEN, H3K4me1 and gain of p-AKT staining, while histologically normal regions maintained PTEN and H3K4me1 staining (**Fig. S2E**). These data indicate that loss of *Kmt2c* or *Kmt2d* alone is insufficient to cause observable histologic dysplasia but cooperates with *Pten* loss in gastric tumorigenesis; they further suggest that *Kmt2d* loss may be more potent than *Kmt2c* loss in promoting gastric cancer initiation.

To examine early direct effects, we focused our analyses on TP, TCD, and TPCD mice at the 3-week and 6-week timepoints. In TP, TCD and TPCD groups, we observed progressively increased thickness of stomach mucosa and increased gross weight of whole stomach (**Figure 2D-F; Figure S3**). Histological analysis revealed mucosal hyperplasia in TP, nuclear dysplasia in TCD, and muscle-invasive gastric cancer in all TPCD stomachs. The cancer exhibits histologic features of Lauren intestinal type [5]. We observed invasion into the submucosa (pT1), the muscularis (pT2), and the serosa (pT3) (**Figure 2G; Figure S4**). Although *Tmprss2-CreER^T2^* can also induces LoxP recombination in epithelial cells of small and large intestines [24], we did not observe any tumors in these tissues at the 6-week timepoint (**Figure 2H**). This is consistent with the low rate of *KMT2C/D* LOF mutations in colorectal adenocarcinoma (**Figure 1C**), indicating that *KMT2C/D* are likely selective tumor suppressors in STAD.

### *Kmt2c/d* knockout impairs gastric lineage differentiation and promotes intestinal metaplasia

The stomach is comprised of gastric glands consisting of pit cells on the surface that form a barrier, stem cells that proliferate and give rise to the other cell types, mucous neck cells that secrete mucus, parietal cells that secrete hydrochloric acid, chief cells that secrete digestive enzymes, enteroendocrine cells that secrete gut hormones, and tuft cells that sense the local environment (**Figure 3A**) [27]. To examine cellular differentiation in GEMMs, we performed IHC and immunofluorescent (IF) staining of gastric lineage markers, including MUC5AC (pit cells), ATP4A (parietal cells), Pepsinogen C (PGC, chief cells and mucous neck cells), and lectin Griffonia simplicifolia-II (GSII, mucous neck cells). In TP mice, we observed mucous neck cell hyperplasia [28], characterized by increased GSII-positive mucous neck cells and decreased ATP4A-positive parietal cells (**Figure 3B**). Ki-67-positive stem/progenitor cells remain in the isthmus region above the mucous neck cells and appear to be slightly expanded (**Figure 3B**). In TCD mice, we observed nuclear dysplasia in the apical region, marked by increased number and proliferation of MUC5AC-positive pit cells (**Figure 3B**) [28]. In TPCD mice, we observed the loss of most gastric lineage makers, along with a disorganized pattern of Ki-67-positive proliferating cells (**Figure 3B**). Compared to normal tissues, the expression of gastric lineage markers was also significantly reduced in human STAD samples (**Figure 3C**).

**Figure 3.**
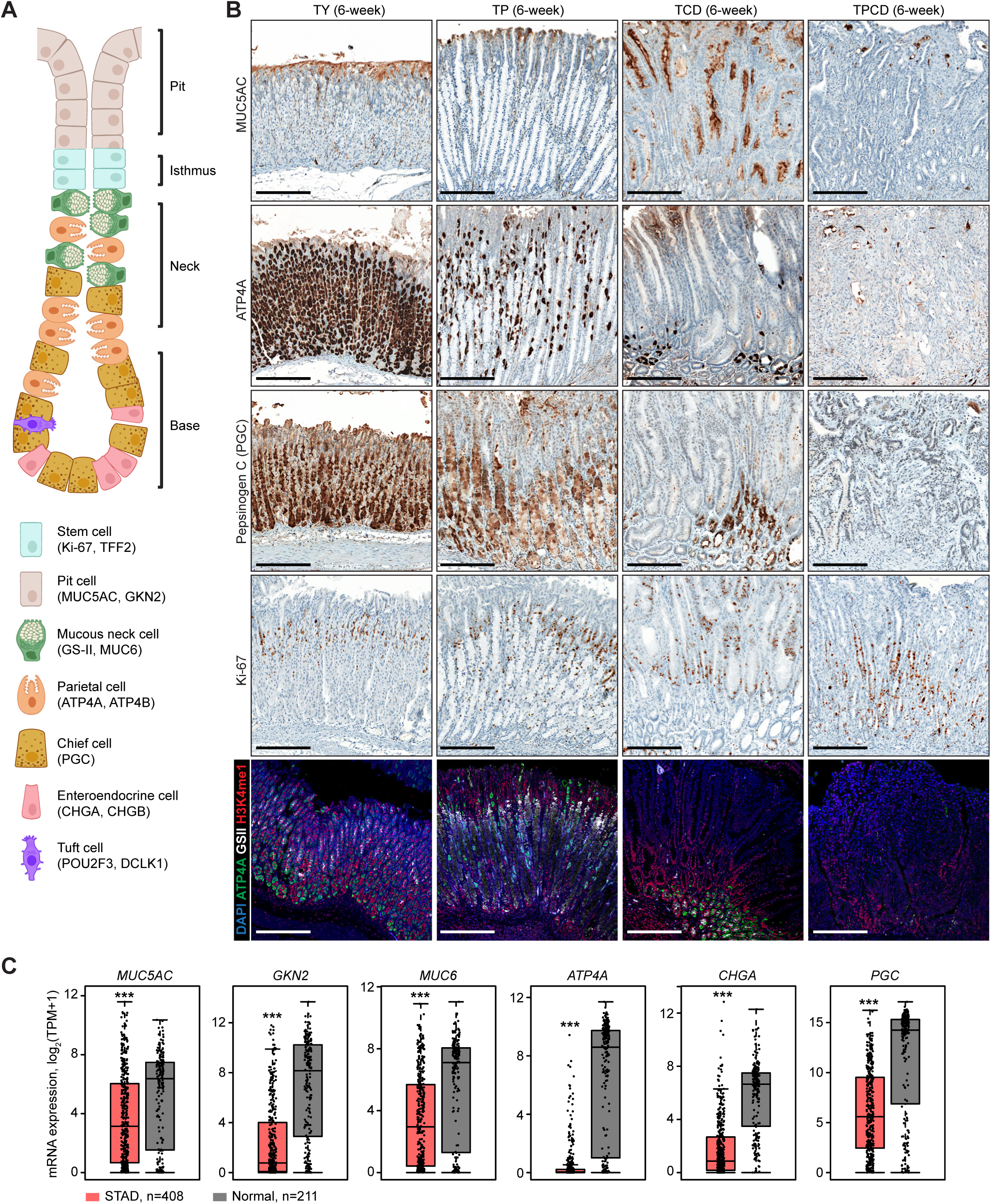
*Kmt2c/d* knockout impairs gastric differentiation. A, Schematic diagram (created with BioRender) illustrating cell types and their distribution in mouse stomach corpus. B, Top, representative immunohistochemistry (IHC) staining of MUC5AC, ATP4A, PGC, and Ki-67 in stomach tissues. Bottom, representative IF of ATP4A, H3K4me1, and lectin GS-II in stomach tissues. Scale bar, 200 µm. C, Expression of *MUC5AC*, *GKN2*, *MUC6*, *ATP4A*, *CHGA*, and *PGC* in STAD (red) and normal stomach (grey) from TCGA dataset. Data were extracted using GEPIA2. The center line represents the median, the box limits represent the upper and lower quartiles, the whiskers represent 1.5 × the interquartile range.

To characterize the diversity of transcriptional alterations, we performed single-cell RNA sequencing (scRNA-seq) on dissociated stomach mucosa 3 weeks post tamoxifen administration (**Figure 4A**). Viable cells were sorted as DAPI-negative using Fluorescence Activated Cell Sorting (FACS) (**Figure S5A**). Cell types were identified using marker genes from prior works and PanglaoDB [27, 29, 30]. We analyzed 47,777 single-cell transcriptomes from TY (n=2), TP (n=3), TCD (n=3), and TPCD (n=2) mice (**Figure S5B**), identifying cell clusters using uniform manifold approximation projection (UMAP) and Leiden clustering.

**Figure 4.**
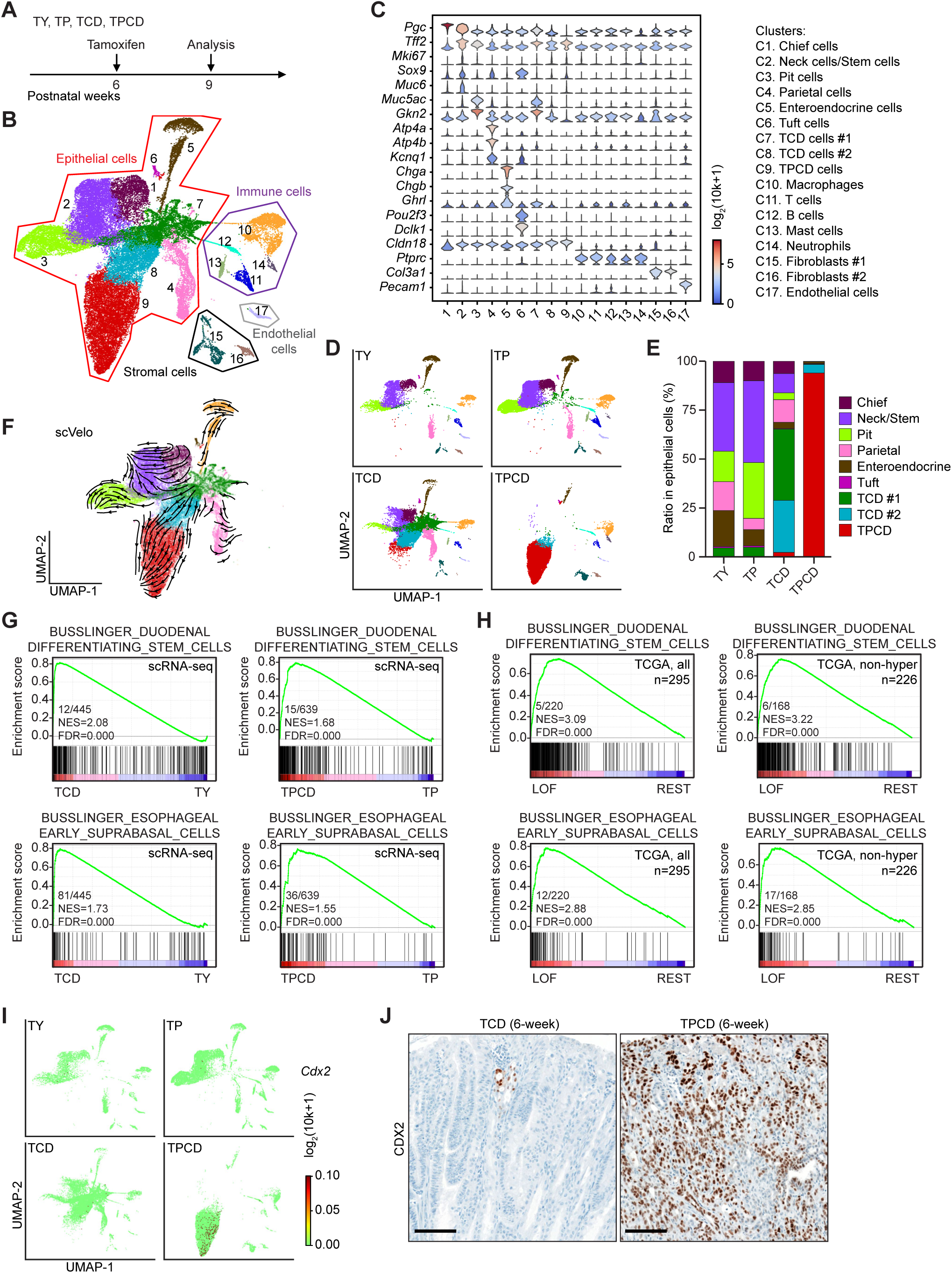
Single-cell RNA-seq reveals impaired differentiation after *Kmt2c/d* loss. A, Schematic of mouse models: two doses of tamoxifen (3 mg × 2) were injected intraperitoneally with a 48-hour interval. Tissues were collected 3 weeks post tamoxifen administration. B, Uniform Manifold Approximation Projection (UMAP) plots showing cell clusters in pooled TY, TP, TCD, and TPCD mice. C, Violin plots of representative cell cluster marker genes. The color in the violin plots indicates the median normalized expression level of genes. D, UMAP plots showing cell clusters in TY, TP, TCD, and TPCD mice. E, Percent of each cell cluster in the pooled epithelial components (C1-C9). F, UMAP-based embedding of RNA velocity analysis showing trajectory transition among cell clusters. G, GSEA analyses in scRNA-seq showing positive enrichment of gene sets associated with duodenal and esophageal lineages following *Kmt2c/d* deletion. H, GSEA analyses in human STAD showing positive enrichment of gene sets associated with duodenal and esophageal lineages in *KMT2C/D*-LOF samples. I, UMAP color-coded by expression of *Cdx2* in TY, TP, TCD, and TPCD cell clusters. J, Representative IHC of CDX2 in stomach tissues of TCD and TPCD mice. Scale bar, 200 µm.

In normal stomachs, we identified clusters of chief cells marked by high *Pgc* expression (C1), neck cells/stem cells marked by high*Tff2*, *Muc6,* and *Mki67* expression (C2), pit cells marked by high *Muc5ac* and *Gkn2* expression (C3) that form a continuum (**Figure 4B-D; Figure S5C, D**). Neck cells and stem cells were placed into the same cluster, but examination of neck cell marker (*Muc6*) and stem cell markers (*Mki67*, *Tff2*) show they occupy different regions of the cluster on UMAP (**Figure S5D**). We further identified parietal cells marked by *Atp4a*, *Atp4b* and *Kcnq1* (C4), enteroendocrine cells marked by *Chga, Chgb* and *Ghrl* (C5), and tuft cells marked by *Pou2f3* and *Dclk1* (C6) (**Figure 4B-D; Figure S5C, D**).

After *Pten* knockout, we observed no new cell clusters but increased ratio of both pit cell and neck/stem cell clusters, consistent with the phenotype of mucous neck cell hyperplasia (**Figure 3B, 4E**). After *Kmt2c/d* knockout, we identified new cell clusters C7 and C8. Cluster C7 exhibited lineage infidelity, with mixed expression of pit cell markers *Muc5ac* and *Gkn2,* neck cell marker *Tff2*, enteroendocrine cell marker *Ghrl* and chief cell marker *Pgc* (**Figure 4C; Figure S5D**), aligning with the phenotype of nuclear dysplasia (**Figure 3B**). Both clusters C7 and C9 exhibited high expression of *Tff2* (**Figure 4C**), a marker of isthmus progenitor cells [31]. In TPCD mice, the dominant cluster C9 showed loss of most gastric epithelial lineage markers (**Figure 4C; Figure S5D**), consistent with the de-differentiation in histological observations.

There was upregulation of many genes upregulated in human stomach cancer, including *Cldn18*, *Onecut2, Cldn4, Trop2, Klf5, Cftr*, and *Plaur*. Interestingly, *Cldn18* was upregulated in cluster C8 and C9 (**Figure 4C; Figure S5D**), suggesting the potential sensitivity of *KMT2C/D*-deficient tumors to Cldn18-based therapies [32]. UMAP-based embedding of RNA velocity analysis revealed the transitional trajectory linking the new clusters C7, C8 and C9 to normal stomach epithelial lineages (**Figure 4F**).

To explore the transcriptional alterations in differentiation, we pooled single-cell transcriptomes of all gastric epithelial cells of each genotype and performed gene set enrichment analysis (GSEA) using cell type signature gene sets (C8) from single-cell sequencing studies of human tissues. We focused on the transcriptional perturbations induced by *Kmt2c/d* knockout. Gene sets associated with duodenal and esophageal differentiation were positively enriched in both TP vs. TY and TPCD vs. TP comparisons (**Figure 4G**), suggesting the induction of dedifferentiation or transdifferentiation following *Kmt2c/d* loss. GSEA of TCGA STAD data also showed positive enrichment of duodenal and esophageal lineage markers in samples with LOF mutations in *KMT2C/D* when we included the entire dataset or when we excluded MSI (non-hyper) samples, suggesting this is not an MSI specific phenomenon (**Figure 4H**). We performed Alcian blue staining of acidic mucin. Alcian blue stains Goblet cells in normal Stomach and is clinically used to identify intestinal metaplasia [28]. We detected Alcian blue positive cells in deep antral gland but not in corpus gland of TY mice (**Figure S6**), consistent with prior reports [28, 33]. In contrast, aberrant Alcian blue-positive staining in supra-basal stomach tissues was observed in TP, TCD, and TPCD mice, reflecting the altered expression of mucins. In TPCD mice, the invasive gastric carcinoma stained positive for Alcian blue. Expression of the intestinal lineage transcription factor CDX2 was only observed in TCD and TPCD mice, suggesting the intestinal differentiation in our models (**Figure 4I, J**). Clinically, CDX2 is found in premalignant gastric lesions that exhibit intestinal metaplasia and in intestinal subtype gastric carcinoma [34]. Together, these data highlight the hyperplastic and metaplastic lineage changes following *Pten* or *Kmt2c/d* loss.

To further investigate the correlation between GEMMs and human gastric cancer development, we conducted integrated analyses of scRNA-seq data with human pre-cancerous and cancerous stomach samples from two recently published datasets [35, 36]. We used the Harmony algorithm to integrate the 3 data sets [37]. Based on the UMAP visualization of the integrated data, we identified four major groups of cell clusters: stromal cells, immune cells, endothelial cells, and epithelial cells (**Figure 5A, B**). Within the endothelial, stromal, and immune clusters, there was good overlap of all the cell populations, providing confidence to the analysis. Here, we focused on the epithelial compartment and included samples of TY, TP, TCD, TPCD, non-atrophic gastritis (NAG), chronic atrophic gastritis (CAG), intestinal metaplasia (IM), early gastric cancer (EGC), diffuse gastric cancer (DGC), intestinal gastric cancer (IGC), and mixed diffuse and intestinal gastric cancer (MixGC) (**Figure 5A, C**). There was consistent overlap of the normal gastric cell types, including the pit cells, parietal cells, neck cells, and chief cells. In TCD mice, there was expansion of Cluster 1 comprised of crypt cell types and extension of few cells into cluster 4 that consists of human gastric intestinal metaplasia and intestinal carcinoma cells (**Figure 5C, D**). In TPCD mice, the vast majority of epithelial cells clustered into cluster 4 (**Figure 5C, D**). These data suggest that the gastric epithelium of TPCD faithfully recapitulates the transcriptional program of intestinal type gastric cancer.

**Figure 5.**
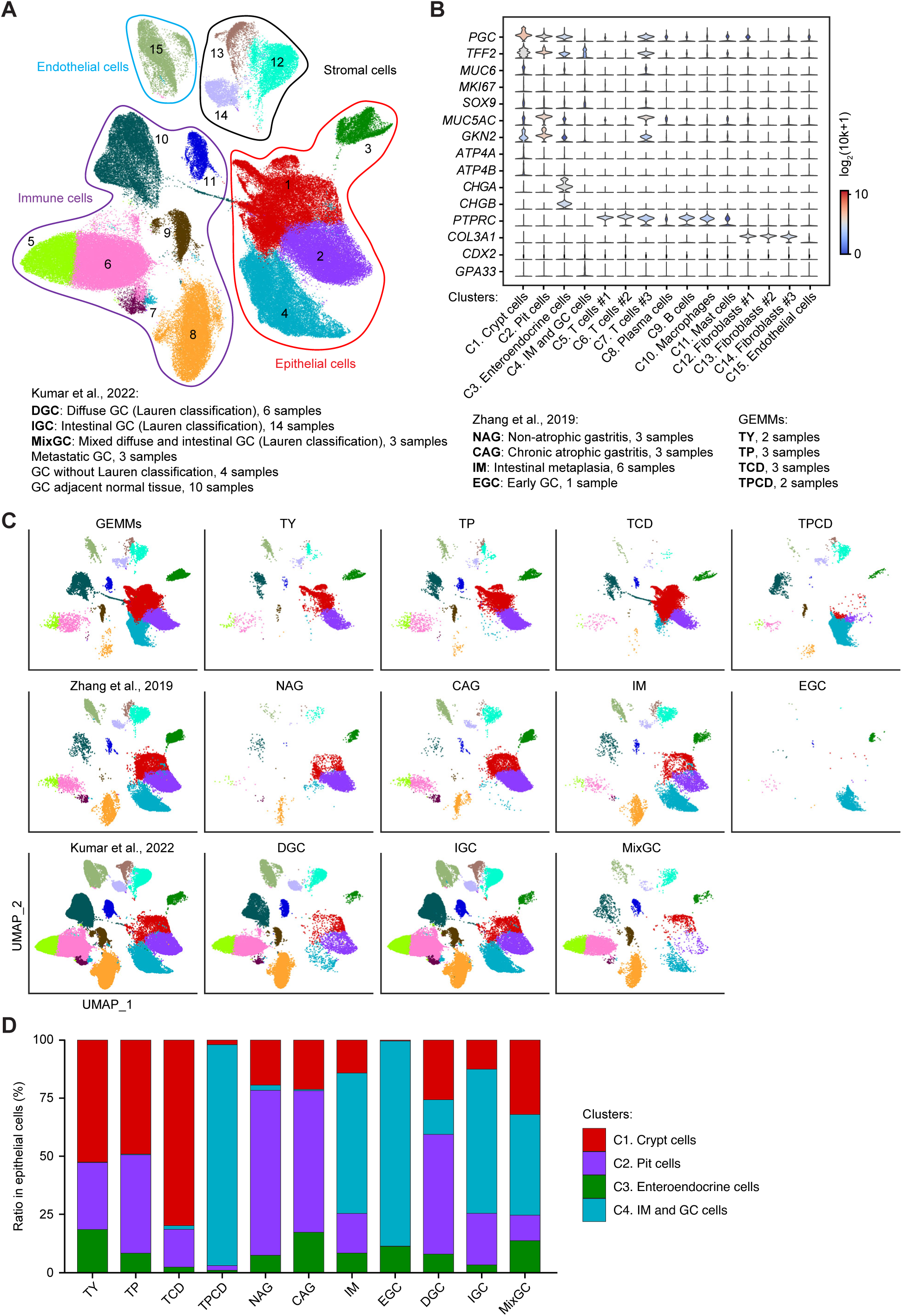
Integrated scRNA-seq analyses reveal the correlation between GEMMs and human GC. A, UMAP plots showing cell clusters of Harmony integrated mouse and two human pre-cancerous and cancerous data sets. B, Violin plots of representative cell cluster marker genes. The color in the violin plots indicates the median normalized expression level of genes. C, UMAP plots showing cell clusters in each subgroup. D, Percent of each cell cluster in the pooled epithelial components (C1–C4).

### *Kmt2c/d* loss enhances MHC-I expression and antigen presentation

Next, we sought to explore potential therapeutic opportunities arising from *Kmt2c/d* loss. GSEA analyses of scRNA-seq data revealed that a gene set associated with antigen presentation was positively enriched after *Kmt2c/d* knockout in both wild-type and *Pten*-loss background (**Figure 6A**). Consistently, comparison of gastric cancers with *KMT2C/D* LOF mutations versus other gastric cancers in TCGA STAD samples also showed positive enrichment of this gene set, even when MSI samples were removed from the analysis (**Figure 6B**). Compared to TY, upregulated expression of MHC-I components *H2-D1*, *H2-K1*, and *B2m* was identified in TP, TCD, and TPCD groups (**Figure 6C**). The upregulated B2M protein level was confirmed by IHC staining (**Figure 6D**; **Figure S7A**). Notably, in TPCD mice with heterogenous *Kmt2c/d* deletion, higher B2M staining intensity was observed in *Kmt2c/d*-deficient neoplastic epithelial cells compared to the adjacent *Kmt2c/d*-intact histologically normal cells (**Figure S7A**), suggesting that the increased B2M level was primarily caused by *Kmt2c/d* deletion rather than the microenvironment.

**Figure 6.**
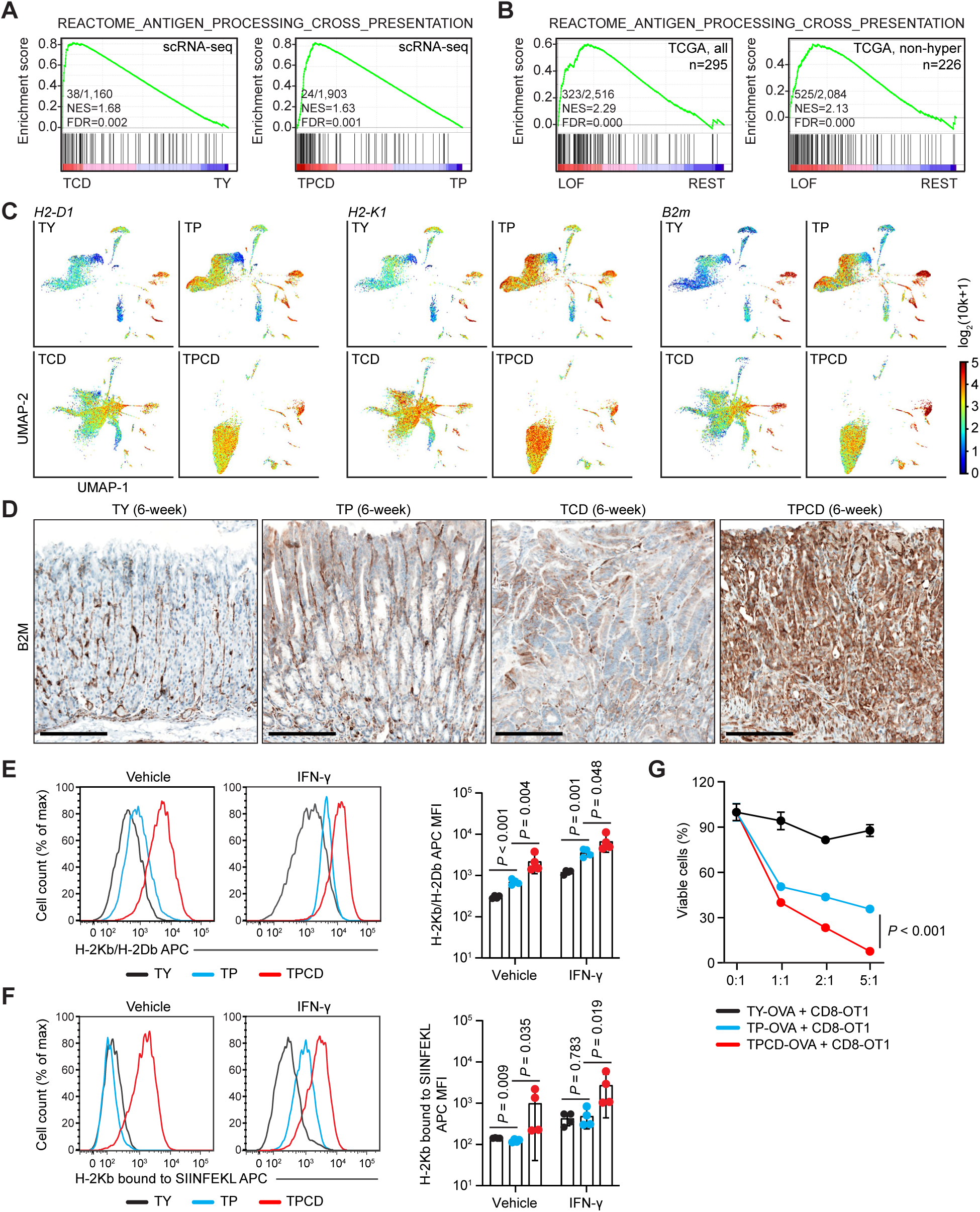
*Kmt2c/d* loss enhances MHC-I expression and antigen presentation. A, GSEA analyses in scRNA-seq showing positive enrichment of gene set associated with antigen presentation following *Kmt2c/d* deletion. B, GSEA analyses in human STAD showing positive enrichment of gene set associated with antigen presentation in *KMT2C/D*-LOF samples. C, UMAP color-coded by expression of *H2-D1*, *H2-K1*, and *B2m* in TY, TP, TCD, and TPCD cell clusters. D, Representative IHC of CDX2 in stomach of TPCD mice. Scale bar, 200 µm. E, Flow cytometry analysis of MHC class I molecules H-2Kb/H2-Db in cultured stomach epithelial cells. To induce the expression of H-2Kb/Db, cells were treated with vehicle or mouse interferon-gamma (IFN-γ, 10 ng/ml) for 24 hours. Data are presented as mean ± SD. Statistical analyses were performed with two-tailed t-test on log_10_ normalized data. F, Flow cytometry analysis of H-2Kb bound SIINFEKL in OVA-expressing stomach epithelial cells. To enhance antigen presentation, cells were treated with vehicle or mouse IFN-γ (10 ng/ml) for 24 hours. Data are presented as mean ± SD. Statistical analyses were performed with two-tailed t-test on log_10_ normalized data. G, Viable OVA-expressing stomach cells after co-culture with OT1 CD8^+^ T cells. Data are presented as mean ± SD. Statistical analysis was performed with two-tailed t-test between TP and TPCD cells in the 5:1 group (T cells : Stomach cells = 5:1).

To assess antigen presentation capability, we generated stomach organoids from TY, TP, and TPCD mice after tamoxifen administration and exogenously expressed chicken ovalbumin (OVA) in these cells. Gastric epithelial cells from TCD mice grew poorly in vitro and could not be studied (**Figure S8A**) We performed PCR using DNA isolated from organoids to confirm efficient deletion of the appropriate alleles (**Figure S8B)**. We used FACS to quantify the expression of MHC-I molecules and the presentation of antigen SIINFEKL in TY, TP and TPCD cells at baseline and after interferon-gamma (IFN-γ) treatment. Consistent with scRNA-seq data, we observed progressively increased cell surfaced expression H2-Db and H2-Kb at baseline level and after IFN-γ stimulation in TP and TPCD cells (**Figure 6E**). Presentation of H2-Kb-bound SIINFEKL, the OVA-derived peptide, was significantly enhanced with *Kmt2c/d* knockout (**Figure 6F**). To functionally assay susceptibility to T-cell mediated, antigen specific killing, we next assayed co-culture of OVA-expressing stomach cells with OT1 CD8^+^ T cells. We observed significantly increased killing in TPCD cells compared to TP cells (**Figure 6G**). These data suggest that *Kmt2c/d* knockout context in gastric cancer is primed for augmented antigen presentation, implying potential sensitivity to immunotherapy, consistent with a prior study in liver cancer [38].

### *Kmt2c/d* loss reduces protein synthesis and confers sensitivity to mTORC1 inhibition

In addition to augmented antigen presentation, we observed that gene sets REACTOME_TRANSLATION and REACTOME_EUKARYOTIC_TRANSLATION_INITIATION were among the most enriched gene sets when comparing TCD vs. TY and TPCD vs. TP in mouse scRNA-seq data (**Figure 7A**). In the TCGA STAD dataset, these gene sets were also enriched in samples with LOF mutations of *KMT2C* and/or *KMT2D* (**Figure 7B**). Interestingly, in *Pten* loss cells that is known to activate mTOR signaling and enhance translation, we found negative enrichment of gene sets associated with protein synthesis (**Figure 7C**). To directly evaluate new protein synthesis, we performed puromycin incorporation assay in the stomachs of TCD mice *in vivo*. Compared to H3K4m1-high cells, reduced puromycin staining intensity was observed in H3K4me1-low cells (**Figure S9A**), indicating decreased new protein synthesis after *Kmt2c/d* loss. We further used O-propargyl-puromycin (OPP), an analog of puromycin, to compare protein synthesis *in vitro* using flow cytometry. Compared to TY cells, *Pten* knockout significantly increased new protein synthesis (**Figure S9B, C**), consistent with prior reports [39]. Further deletion of *Kmt2c/d* reduced protein synthesis in TPCD compared to TP cells (**Figure S9B, C**), indicating differential function of *Kmt2c/d* and *Pten* in regulating new protein synthesis. These data indicate that in the gastric epithelium, there is an inverse correlation between the mRNA expression of ribosomal protein genes and the overall rate of protein translation and posit that upregulation of gene sets associated with translation upon *Kmt2c/d* loss, mostly comprised of ribosomal proteins, may be a response to inadequate translation [40].

**Figure 7.**
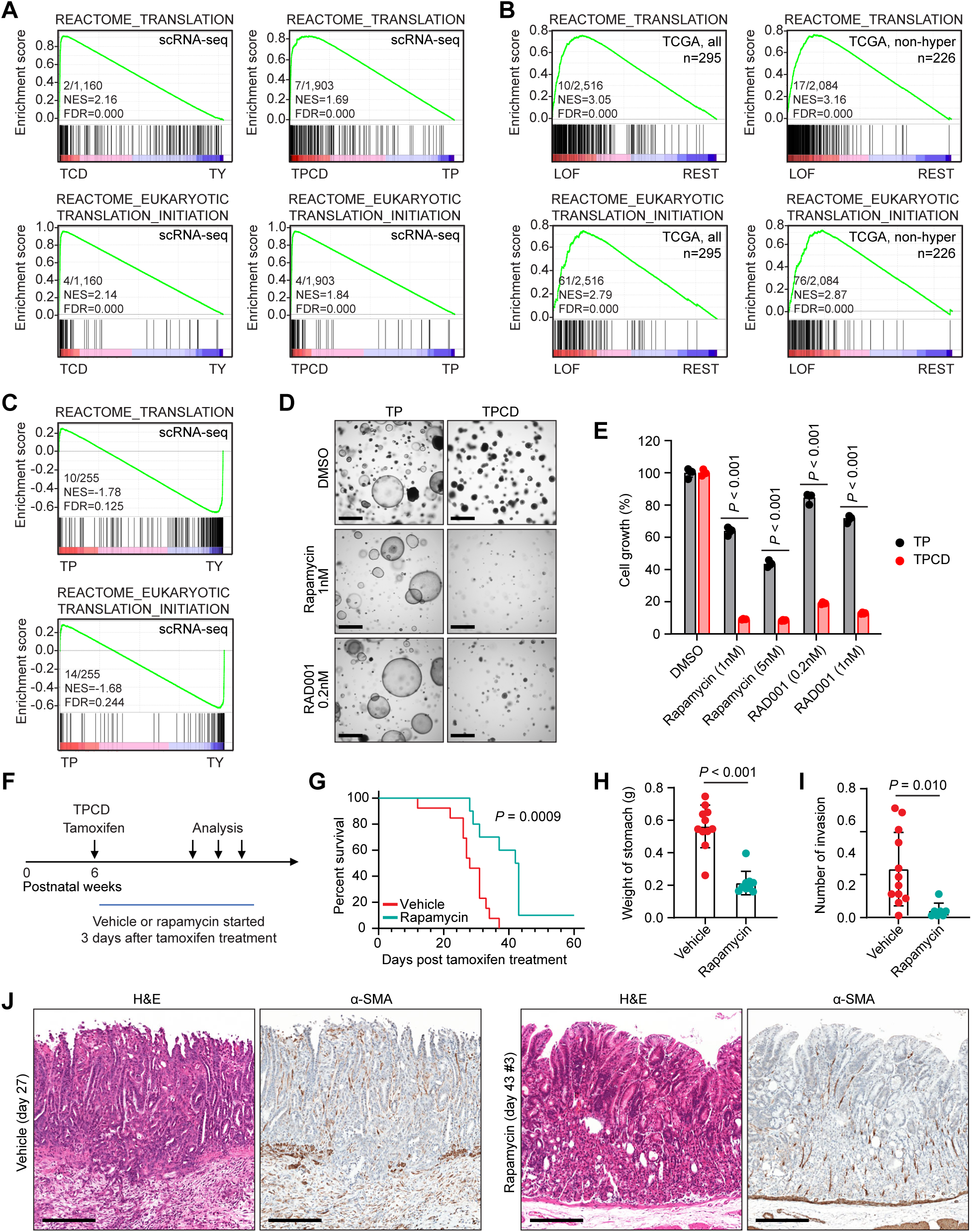
*Kmt2c/d* loss confers sensitivity to mTORC1 inhibition. dA, GSEA analyses in scRNA-seq showing positive enrichment of gene sets associated with protein translation following *Kmt2c/d* deletion. B, GSEA analyses in human STAD showing positive enrichment of gene sets associated with protein translation in *KMT2C/D*-LOF samples. C, GSEA analyses in scRNA-seq showing negative enrichment of gene sets associated with protein translation following *Pten* deletion. D-E, Representative bright-field images of organoid from TP and TPCD groups. Scale bar, 1mm. Cells were seeded in Matrigel (500 cells per blob, 50µl), and treated from day 2 for 10 days. Organoid culture medium and inhibitors were refreshed every 4 days. Cell growth was measured using the CellTiter-Glo luminescent reagent. Data are presented as mean ± SD and analyzed with two-tailed t-test. F, Schematic illustration showing the induction and treatment of stomach cancer in TPCD mice. Vehicle or rapamycin treatment started 3 days after the first dose of tamoxifen. G, Kaplan-Meier plots showing the survival of mice treated with vehicle or rapamycin. H, Stomach weight in vehicle- or rapamycin-treated TPCD mice. Data are presented as mean ± SD and analyzed with two-tailed t-test. I, Statistics of invasive lesions in vehicle- or rapamycin-treated TPCD stomach. Invasion sites were determined using IHC of alpha-smooth muscle actin (α-SMA). Data are presented as mean ± SD and analyzed with two-tailed t-test. J, Representative H&E and IHC of α-SMA in vehicle- or rapamycin-treated TPCD mice. Scale bar, 200 µm.

Our data indicate that *Kmt2c/d* loss leads to compromised protein translation, and we tested whether this may lead to sensitivity to protein synthesis inhibitors. We performed cell viability assay using mTORC1-specific inhibitors (rapamycin and RAD001) and the dual mTORC1/2 inhibitor INK-128. TPCD cells were more sensitive to these inhibitors than TP cells, with greater difference with mTORC1-specific inhibitors (**Figure S9D, E**). Increased sensitivity to mTORC1 inhibitors was also observed in TPCD organoids (**Figure 7D, E**).

To determine if rapamycin can affect gastric tumorigenesis in TPCD GEMM mice, we treated TPCD mice with vehicle or rapamycin starting three days post tamoxifen administration (**Figure 7F**). Rapamycin treatment significantly prolonged the survival of mice and reduced gross stomach weight (**Figure 7G, H**). Histological staining showed reduced submucosal invasion and improved gastric glandular organization appreciated from IHC of α-SMA, GKN2, and PGC (**Figure 6I, J; Figure S10, S11**). Together, these results suggest that *Kmt2c/d* loss confers sensitivity to mTORC1 inhibition.

### Combination of rapamycin and anti-PD1 suppresses TPCD cell growth *in vivo*

To evaluate the combinatorial effects of mTORC1 inhibition and immunotherapy *in vivo*, we grafted TPCD tumor cells into C57BL/6 mice and treated them with vehicle, rapamycin, anti-PD1 or the combination. Monotherapy with either rapamycin or anti-PD1 effectively suppressed tumor growth, reduced tumor formation rate from 8/8 (vehicle) to 6/8 (anti-PD1) and 3/8 (rapamycin) (**Figure 8A, B**). The combination further inhibited tumorigenesis, with a 1/8 tumor formation rate by day 28 pos-grafting, though no obvious synergy was observed (**Figure 8B**). Consistent with the secretion function of stomach mucosa [41], fluid secretion was observed in some tumors in the control group (**Figure 8A**). After puncturing the tumors and aspirating the fluid, a significant reduction in tumor weight was identified (**Figure 8C**). Treatment with rapamycin led to decreased p-S6 (Ser235/236) level, and anti-PD1 treatment increased CD8^+^ T cell infiltration, confirming the efficacy of these treatments (**Figure 8D**). Additionally, rapamycin treatment improved histological differentiation, although no change in CDX2 expression was observed (**Figure 8D**). Together, our findings highlight the potential of combining mTORC1 inhibitors and immune checkpoint blockade for *KMT2C/D*-deficient STAD treatment.

**Figure 8.**
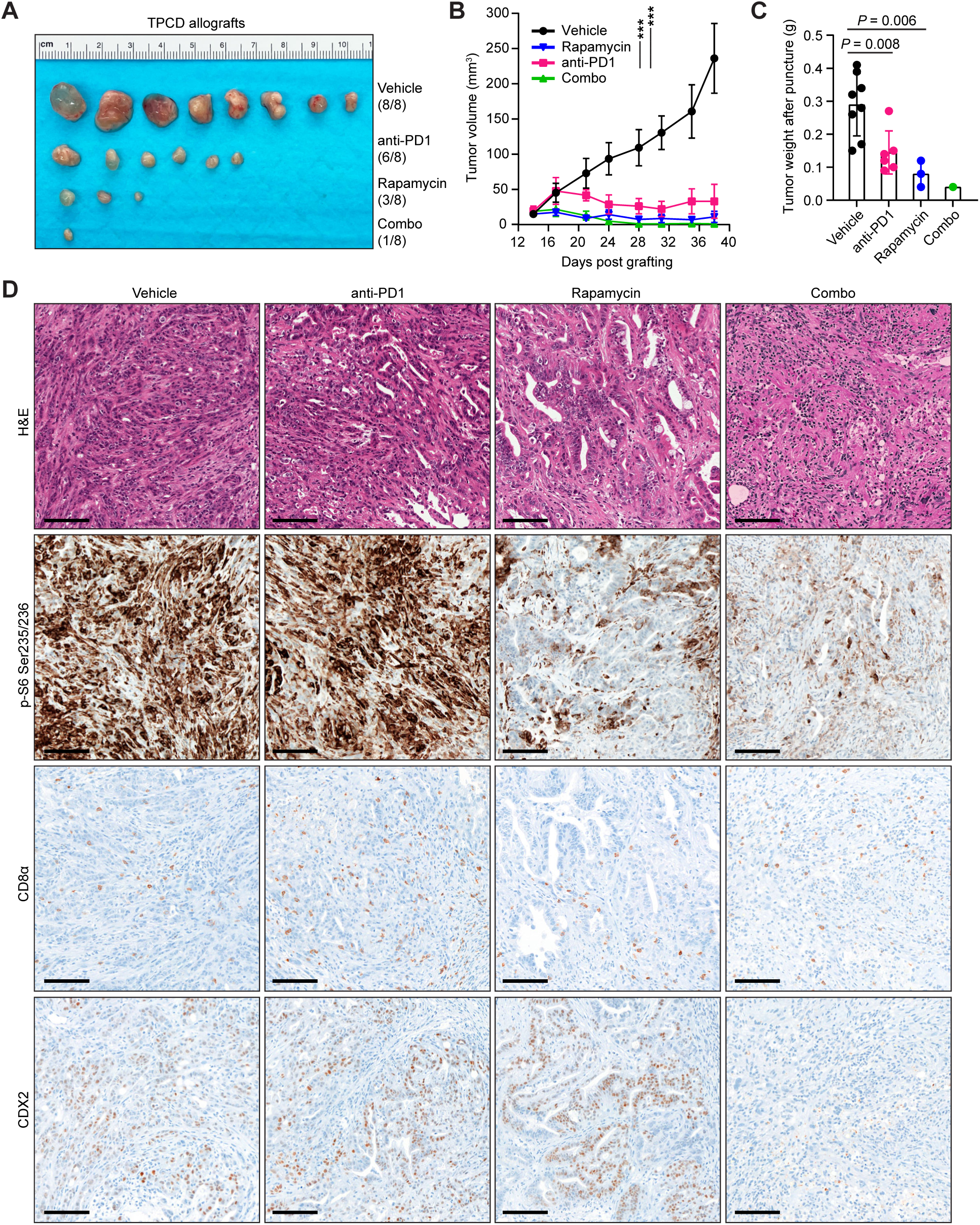
Combination of rapamycin and anti-PD1 suppresses growth of TPCD cells *in vivo*. A-B, Allografts and growth curves of TPCD tumors treated with rapamycin (5 mg/kg/day) or vehicle control in C57BL/6 mice. Treatment started 2 weeks after injection of cells into the mammary fat pad. Data are presented as mean ± SEM and analyzed with two-tailed t-test at endpoint. C, Statistics of TPCD tumor weight in C57BL/6 mice. Data are presented as mean ± SD and analyzed with two-tailed t-test. D, Representative H&E and IHC staining of p-S6 Ser235/236, CD8α, and CDX2 in grafted TPCD tumor tissues. Scale bar, 100 µm.

## Discussion

The identification of driver mutations in MSI tumors is difficult due to large number of passenger mutations, particularly in large genes. The TCGA specifically excluded MSI tumors from analysis of significantly mutated genes [3, 20, 21] However, hotspot mutations in oncogenes are well known drivers in MSI cancer, such as *KRAS, PIK3CA,* and *BRAF^V600E^*. A comparison of KMT-family of histone methyltransferase genes between gastric and colorectal cancer found *KMT2C* to be specifically mutated in MSI gastric cancers and mutated samples lost all protein staining by IHC [42]. A pan-TCGA analysis of microsatellite instability that specifically analyzed frame-shift events at microsatellites found cancer-specific preferences, including higher frequency of events at microsatellites in *KMT2C* and *KMT2D* in gastric cancer [43]. Our analysis of TCGA data suggests the positive selection of *KMT2C* and *KMT2D* LOF mutations in STAD, as *KMT2C* and *KMT2D* LOF are much less common in MSI cancers in other cancer types and other similarly large genes do not harbor as many LOF mutations in STAD.

These data led us to study the in vivo role of *Kmt2c* and *Kmt2d* in GEMMs. Our data showed that *Kmt2c/d* loss promotes gastric cancer initiation and drives significant molecular and phenotypic changes, including impaired cellular differentiation, enhanced antigen presentation, and reduced protein synthesis. Notably, in the small and large intestines, these changes were not observed, recapitulating human pathology. These findings provide critical insights into the role of *Kmt2c/d* as key regulators of gastric epithelial homeostasis and tumorigenesis. Despite the high prevalence of *KMT2C/D* LOF mutations in STAD, knockout of *Kmt2c* and/or *Kmt2d* alone was insufficient to induce cancer initiation. A secondary oncogenic mutation, such as *Pten* loss, is required to fully drive tumorigenesis, consistent with reports in lung cancer and urothelial cancer [10, 12]. The inability of *Kmt2c/d* loss alone to initiate cancer may be attributed to the decreased new protein synthesis, implicating the potential tumor-promoting role of PI3K and MAPK signaling in the context of *Kmt2c/d* deficiency.

Precancerous lesions are considered precursors in gastric cancer initiation [44, 45]. Consistent with scRNA-seq data from human stomach intestinal metaplasia tissues [44], loss of *Kmt2c/d* led to dysplasia and expansion cells with mixed pit, neck, and stem cell features. These observations were accompanied by the mosaic expression of alcian blue-positive mucin and positive staining of CDX2 in TCD mice, suggesting the emergence of intestinal metaplasia and the high relevance of our GEMMs to clinical observations. Notably, recent sequencing studies have identified *KMT2C* and *KMT2D* nonsynonymous mutations in intestinal metaplasia, and further identified *KMT2D* as one of the driver genes in intestinal metaplasia progression [45, 46]. In the TPCD stomach cancer model, we observed significantly reduced expression of key gastric lineage markers, indicating a shift toward a less differentiated state. However, this model expresses high levels of Claudin-18, an FDA approved therapeutic target in gastric cancer [47, 48], and may be a useful model for mechanistic and preclinical studies. These molecular and histological alterations closely mirror the progression of human STAD, reinforcing the relevance of our findings to human disease.

Our data reveal upregulated MHC-I molecule expression and enhanced antigen presentation in *Kmt2c/d*-deficient cells, suggesting a potential vulnerability to immune-based therapies. This is further supported by the increased sensitivity of *Kmt2c/d*-deficient cells to OT1 CD8^+^ T cells *in vitro* and to anti-PD1 treatment *in vivo*, consistent with observations in liver and bladder cancer [10, 38]. Our data is consistent with clinical observations of improved response to immune checkpoint blockade in tumors with *KMT2C* or *KMT2D* mutations [10, 49]. We acknowledge that checkpoint blockade is already approved for most gastric cancers and is highly active in the MSI subgroup [50], which may limit the clinical utility of this finding for anti-PD1 monotherapy. We further found that loss of *Kmt2c/d* led to decreased protein translation both *in vivo* and in organoids, leading to sensitization to mTORC1 inhibitors in both organoids *in vitro* and allograft tumors *in vivo*. Everolimus has been previously studied in a phase 3 trial in previously treated advanced gastric cancer. In the overall population, there was a small but significant improvement in progression free survival but only a trend for overall survival [51]. Our data suggests that loss of *KMT2C/D* may be a biomarker of those who benefit. Our findings highlight the potential of combining mTORC1 inhibitors with immune checkpoint blockade as a therapeutic strategy for *KMT2C/D*-deficient STAD. Rapamycin is a potent immunosuppressant and combination with anti-PD1 may seem counterintuitive. However, several recent studies have shown that rapamycin and JAK inhibitors paradoxically improve efficacy of anti-PD1 therapy [52, 53]. Moreover, rapamycin has been shown to suppress growth of *Arid1a*-deficient stomach cancer cells [7], suggesting a broader applicability of combining mTORC1 inhibitors and immune checkpoint blockade in STAD treatment.

Our study has several limitations. *Tmprss2-CreER^T2^* is active in several endoderm-derived epithelial lineages and tamoxifen injection induces robust recombination in the stomach, intestines, bladder, and luminal prostate epithelial cells. This limits long-term studies and the ability to evaluate metastasis in the GEMM. We have used organoids and allografts from GEMM mice to orthogonally validate our findings. Methods to localize recombination may overcome some limitations [23].

## Conflict of interests

P.C. has received personal honoraria/advisory boards/consulting fees from Deciphera, Exelixis, Zai Lab, Novartis, Ningbo NewBay Medical Technology; P.C. has received institutional research funding from Pfizer/Array, Novartis, Deciphera, Ningbo NewBay Medical Technology. Y.C. has stock ownership and received royalties from Oric Pharmaceuticals. Y. J. has received personal advisory boards/consulting fees from Abbvie, Alphasights, Amerisource Bergen, Ask-Gene Pharma, Inc., Arcus Biosciences, Astellas, Astra Zeneca, Basilea Pharmaceutica, Bayer, Boehringer Ingelheim, Bristol-Myers Squibb, Clinical Care Options, Daiichi-Sankyo, eChina Health, Ed Med, Resources (OncInfo), Eisai, Eli Lilly, Geneos Therapeutics, GlaxoSmithKline, Guardant Health, Inc., H.C. Wainwright & Co., Health Advances, HMP Global, Imedex, Imugene, Inspirna, Lynx Health, Mashup Media LLC, Master Clinician Alliance, Merck, Merck, Serono, Mersana Therapeutics, Michael J. Hennessy Associates, Oncology News, Paradigm Medical Communications, PeerMD, PeerView Institute, Pfizer, Physician’s Education Resource, LLC, Research to Practice, Sanofi Genzyme, Seagen, Silverback Therapeutics, Suzhou Liangyihui Network Technology Co., Ltd, Talem Health, TotalCME, Zymeworks Inc. Y. J. has stock options from Inspirna and Veda Life Sciences, Inc. The other authors declare no competing interests related to this study.

## Author contributions

Conceptualization: N. W., Y. C.

Investigation: N. W., T. Z., M. R. P., W. H. C., M. N. K., D. M. S

Formal analysis: D. L., N. W., Y. C.

Histology review: Y. B., L. T., Y. J.

Supervision: Y. C., C. P.

Writing: N. W., D. L., T. Z., M. L. Y. J. P. C., Y. C.

## Acknowledgements

This work was supported by grants from the National Institute of Health (NIH) and National Cancer Institute (NCI) grants (R01CA228216, DP2CA174499, P50CA217694) to P.C.; (U54CA224079, P50CA092629, P50CA221745, R01CA193837, U01CA224044, R01CA208100) to Y.C.; Samuel Waxman Cancer Research Foundation to Y.C; Gladstein Family Bladder Cancer Research Fund to Y.C.; Geoffrey Beene Cancer Research Fund to P.C., Y.C.; and Department of Defense (DOD) grant (W81XWH-15-1-0124), Francis Collins Scholar NTAP, Cycle for Survival and Linn Family Discovery Fund to P.C. We thank the following core facilities at MSKCC: Flow Cytometry, Integrated Genomics Operations, Molecular Cytology, and Research Animal Resource Center.

## Methods and Materials

### Mouse studies

Mouse experiments were approved by the Institutional Animal Care and Use Committee of Memorial Sloan Kettering Cancer Center, New York. Mice were maintained under specific-pathogen-free conditions, with a 12-hour light/dark cycle (lights on/off at 6am/pm), controlled temperature (18-24℃) and humidity (40-60%), and access to chow and sterilized water *ad libitum*. The following genetically engineered mouse strains were used: *Tmprss2-CreER^T2^-IRES-nlsEGFP* (*Tmprss2^tm1.1(cre/ERT2)Ychen^*, MGI:5911389), *Pten^flox^*(*Pten^tm2.1Ppp^*, MGI:2679886), *Kmt2c^flox^* (Exon 3 flanked by LoxP sites), *Kmt2d^flox^* (Exon 50-51 flanked by LoxP sites), and *Rosa26-CAG-LSL-EYFP* (*B6.Cg-Gt(ROSA)26Sor^tm3(CAG-EYFP)Hze^*, stock no. 007903, Jackson Laboratory) [10]. To induce *Tmprss2-CreER^T2^* activity, two doses of tamoxifen (Toronto Research Chemicals, T006000, 3 mg per dose in corn oil) were administered intraperitoneally to 8-12 week-old mice with interval of 48 hours.

To test the effect of rapamycin on stomach cancer progression in GEMMs, we administered tamoxifen to *Tmprss2-CreER^T2^;Pten^flf^;Kmt2c^f/f^;Kmt2d^f/^* (TPCD) mice to induce gene knockout. Three days after the first dose of tamoxifen administration, mice were treated with either vehicle (5% ethanol, 5% PEG-400, 5% Tween-80 in PBS, 200 µL per injection) or rapamycin (#HY-10219, Med Chem Express, 5 mg/kg, once daily, 5 days a week, 200 µL per injection) via intraperitoneal injection.

To assess the *in vivo* responses to rapamycin and immune checkpoint blockade, we grafted 5 million TPCD cells into the mammary fat pad of 6-8-week-old female mice (C57BL/6J, stock no. 000664, Jackson Laboratory). Two weeks after grafting, mice were treated with vehicle, rapamycin, IgG control (BE0089, Bio X Cell, 8mg/kg, twice a week, 200 µL in PBS), or anti-PD1 (BE0146, Bio X Cell, 8 mg/kg, twice a week, 200 µL per injection). Tumor sizes were measured twice a week using a digital caliber and calculated using the formula 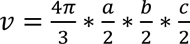, where a, b, and c represent the length, width, and thickness of the tumor, respectively.

### scRNA-seq and analysis

Stomachs were collected from TY (n=2 mice), TP (n=3 mice), TCD (n=3 mice), and TPCD (n=2 mice) groups three weeks after tamoxifen administration. Stomach mucosae were stripped using tweezers and washed with cold PBS. After mincing with a scalpel, the mucosal tissues were digested for 30 minutes with TrypLE (12605010, Gibco), followed by 45 minutes with collagenase/hyaluronidase (07912, STEMCELL Technologies). Viable cells were sorted as DAPI-negative using a BD FACSymphony^TM^ S6 Cell Sorter. For each mouse, 10,000 cells were processed for encapsulation and library preparation (Chromium Next GEM Single Cell 3’ GEM, Library and Gel Bead Kit, 10X Genomics). For each sample, 200 million reads were acquired on a NovaSeq S4 flow cell platform.

Single-cell RNA-seq data were analyzed as previously described [10, 54]. Briefly, sequencing reads were mapped to the mouse genome (GRCm38) using the Cell Ranger (7.0.0) software (10X Genomics). Downstream analysis and figure plotting were processed using Scanpy (1.9.8) [55]. Cells were removed if they expressed fewer than 100 unique genes, fewer than 2,000 total counts, more than 40,000 total counts, or greater than 20% mitochondrial reads. Genes detected in less than 20 cells and all mitochondrial genes were excluded from subsequent analyses. Combining samples from all cohorts yielded a count matrix of 47,777 cells by 20,618 genes, with a median of 9,037 counts and a median of 2,575 genes per cell. The count matrix was normalized by log_2_(10K+1) to identify the top 2,000 highly-variable genes. The count matrix was further scaled to a mean of 0 and a standard deviation of 1 for principal component analysis (PCA), UMAP dimensionality reduction, and Leiden clustering [56]. PCA was performed on the 1,000 most variable genes and the top 50 principal components retained 41% of the variance. Cell types were determined using a combination of markers genes identified in prior literature and the web-based tool Panglao DB (https://panglaodb.se/) [29].

Differentially expressed genes among each group were compared using Scanpy (1.9.8) [55]. For GSEA comparing genotypes of gastric cells using mouse scRNA-seq data, we pooled all gastric epithelial cells of each genotype to generate a pseudo-bulk expression data. We generated the ranked ordered list using the difference of pseudo-bulk Log2 expression between genotypes. For GSEA comparing *KMT2C/D* LOF samples and other samples, we calculated the mean Log_2_ expression of all genes in samples with *KMT2C* or *KMT2D* LOF mutations (nonsense or splice site) and in the remaining samples. We generated the ranked list using the difference of mean Log_2_ expressions. GSEA was performed on the ranked gene list using the JAVA GSEA 4.1.0 program, with curated gene sets (C2, C8) and the Hallmark gene set (H) from the Molecular Signatures Database v7.4 using gene set permutation.

The processed scRNA-seq data from human precancerous and cancerous stomachs were downloaded from GSE134520 and GSE183904. We used the same filtration parameters as the original studies. Harmony (0.0.10) algorithm was used for the integrated analyses [37].

### Mouse stomach cell culture

Mouse stomach epithelial cells were isolated as described above. Cells were cultured in growth factor reduced Matrigel matrix (356231, Corning) using organoid culture medium. The medium formulation is as follows: advanced DMEM/F12 (12634010, Gibo), B27 supplement (2% v/v, 17504044, Gibco), Fetal Bovine Serum (5% v/v, FB-11, Omega Scientific), Noggin conditioned medium (10% v/v), R-Spondin conditioned medium (10% v/v), Wnt-3A conditioned medium (50%, v/v), HEPES (2mM, pH 7.4), EGF (50 ng/ml, AF-100-15, PeproTech), FGF2 (200 ng/ml, AF-100-26, PeproTech), Y-27632 (10µM, S1049, Selleck Chemicals), A83-01 (0.5µM, S7692, Selleckchem), N-acetyl-L-cysteine (1.25mM, A9165, Millipore Sigma), SB202190 (10µM, S1077, Selleck Chemicals), Nicotinamide (10mM, N0636, Millipore Sigma), Gastrin (10nM, G9145, Sigma), Primocin (100 µg/ml, ant-pm-2, InvivoGen), penicillin-streptomycin (1% v/v, 15140122, Gibco), L-glutamine (1% v/v, 25030081, Gibco), and GlutaMAX (1% v/v, 35050061, Gibco). Successful deletions were validated using primers amplifying the floxed alleles. Primers for genotyping are listed in Table S1.

To assess cellular responses to mTORC1 and mTORC2 inhibitors in Matrigel culture, TP and TPCD cells were mixed 1:2 with Matrigel and seeded at 500 cells per blob (50 µL final volume). Cells were treated starting from day 2 for 10 days, with the medium and inhibitors refreshed every four days. At the endpoint, images of organoid were captured using a Nikon ECLIPSE Ti2 inverted microscope. Organoids were digested with TrypLE for 30 minutes at 37℃. After centrifugation, cells were lysed with CellTiter-Glo luminescent reagent (G9243, Promega) and measured using the GLOMAX 96-microplate luminometer (Promega).

To culture stomach epithelial cells under two-dimensional conditions, plates or dishes were coated with collagen I (A1048301, Gibco, 1:100 dilution in cold-sterilized water) for 1 hour at room temperature [27]. Cells were cultured using the same stomach organoid medium. To test dose responses to mTORC1 and mTORC2 inhibitors (Rapamycin, S1039; RAD001, S1120; INK-128, S2811; Selleck Chemicals), 1,000 TP and TPCD cells were seeded in pre-coated 96-well plate (100µL final volume). Cells were treated starting from day 2 for 5 days, and cell viability was measured using the CellTiter-Glo luminescent reagent on a GLOMAX 96-microplate luminometer

### OVA antigen presentation and OT1 CD8^+^ T cell killing assay

OT1 CD8^+^ T cells were negatively isolated by depletion of magnetically labelled cells (130-104-075, Miltenyi Biotec) from the spleen of OT-1 mice (C57BL/6-Tg(TcraTcrb)1100Mjb/J, 003831, Jackson Laboratory). CD8^+^ T cells Plates were pre-coated with anti-CD3 antibody (2 µg/ml in PBS, 100302, BioLegend) overnight at 4℃. Naïve CD8+ T cells were culture in plate-bound anti-CD3 antibody and soluble anti-CD28 (2 µg/ml, 102102, BioLegend) antibody for 48 hours [57]. T cell culture medium was prepared using RPMI-1640, supplemented with fetal bovine serum (10% v/v), penicillin-streptomycin (1% v/v), L-glutamine (1% v/v), 2-Mercaptoethanol (1:1000, 21985-023, Gibco), and interleukin 2 (10 ng/ml, 402-ML, R&D Systems). CD8^+^ T cells were then expanded for 2 more days in T cell culture medium.

To induce presentation OT-1-binding antigen SIINFEKL in stomach cells, we used the lentiviral vector pLVX-puro-cOVA (135073, Addgene) to exogenously express chicken ovalbumin protein. We detected the expression of MHC-I components (H-2Kb/Db-APC, 114614, BioLegend) and the presentation of SIINFEKL (H-2Kb bound SIINFEKL-APC, 141606, BioLegend) using a BD LSRFortessa instrument. In the co-culture killing assay, 50,000 TY-OVA, TP-OVA, and TPCD-OVA cells were seeded in collagen I-coated 12-well plates. On the second day, medium was replaced with T cells culture medium containing mouse interferon-gamma (10 ng/ml, 485-MI, R&D Systems). Twenty-four hours later, OT1 CD8^+^ T cells were mixed with tumor cells at ratios of 1: 1, 2:1, and 5:1. After overnight incubation, floating T cells and dead stomach cells were decanted and washed twice with PBS. The remaining cells were measured using CellTiter-Glo luminescent reagent on a GLOMAX 96-microplate luminometer.

### Histological analysis

Mouse stomach, intestine, and colon tissues were dissected and opened longitudinally. After washing in cold PBS, tissues were fixed in 4% paraformaldehyde, dehydrated, and embedded in paraffin. Paraffin embedding and sectioning was performed by Histoserv Inc. IHC was conducted using a Ventana automatic stainer. The following primary antibodies were used in this study: H3K4me1 (5326, Cell Signaling Technology, 1:100), Pepsinogen C (PGC, ab180709, Abcam,1:2,000), MUC5AC (MA1-21907, Thermo Fisher Scientific, 1:1,000), ATP4A (D031-3, MBL Life Science, 1:1,000), Ki-67 (ab16667, Abcam, 1:100), CDX2 (MA5-14494, Thermo Fisher Scientific, 1:500), alpha-smooth muscle actin (α-SMA, ab5694, Abcam, 1:1,000), B2M (HPA006361, Millipore Sigma, 1:500), CD8 alpha (98941, Cell Signaling Technology, 1:100), p-S6 Ser235/236 (2211, Cell Signaling Technology, 1:200), PTEN (9188, Cell Signaling Technology, 1:100), and p-AKT Ser473 (4060, Cell Signaling Technology, 1:100). Both H&E and IHC slides were scanned using a Mirax digital slide scanner. The thickness of stomach mucosa was measured using QuPath-0.4.4.

Immunofluorescent staining was performed using primary antibodies against EYFP (2956, Cell Signaling Technology, 1:100), H3K4me1 (5326, Cell Signaling Technology, 1:100), ATP4A (D031-3, MBL Life Science, 1:500), and Puromycin (MABE343, Millipore Sigma, 1:100). Secondary antibodies conjugated with Alexa Fluor 488 (A11008, Thermo Fisher Scientific, 1:400) or Alexa Fluor 555 (A21424, Thermo Fisher Scientific, 1:400) were used in this study. The lectin GS-II Alexa Fluor 647 (L32451, Thermo Fisher Scientific, 1:500) was applied along with secondary antibodies. Fluorescent images were captured using a Leica TCS SP5 inverted confocal microscope.

Alcian blue staining was performed using paraffin sections. The alcian blue staining solution was prepared by dissolving 1 gram of alcian blue (8GX) (A5268, Millipore Sigma) in 100 ml of 3% acetic acid at pH 2.5. Nuclear fast red staining solution was prepared by dissolving 1 gram of nuclear fast red (60700, Millipore Sigma) and 50 grams of aluminum sulfate (227617, Millipore Sigma) in 1 litter of water. After dewaxing, sections were permeabilized with 0.5% Triton-X 100 for 10 minutes, stained with alcian blue solution for 30 minutes, and counterstained with nuclear fast red solution for 5 minutes. Images were scanned using a Mirax digital slide scanner.

BaseScope was performed using Leica Bond RX as previously described [10]. Fresh sections were stained with probes for *Kmt2c* (#1285828-C1, ACD Bio) and *Kmt2d* (#1285848-C1, ACD Bio). BaseScope™ LS Reagent Kit-RED was used to visualize the signal (#323600, ACD Bio). Sections were scanned using Mirax Digital Slide scanner.

### Human stomach adenocarcinoma dataset analysis

To analyze gene expression differences between *KMT2C/D*-TRUNC samples and the remaining samples, we used the cBioPortal annotation of the TCGA STAD (https://www.cbioportal.org/study/summary?id=stad_tcga_pub), Colorectal (https://www.cbioportal.org/study/summary?id=coadread_tcga_pub) and Endometrial datasets (https://www.cbioportal.org/study/summary?id=ucec_tcga_pub). Analysis and OncoPrint were performed on cBioPortal. Comparisons were made using all samples (n=295) or non-hypermutated samples (n=226). GSEA was performed on the ranked gene list using the JAVA GSEA 4.1.0 program, with curated gene sets (C2, C8) and the Hallmark gene set (H) from the Molecular Signatures Database v7.4.

### Western blot

Stomach epithelial cells were cultured under two-dimensional condition, and treated with DMSO, 1 nM Rapamycin (S1039, Selleck Chemicals), and 5 nM Rapamycin for 1 hour. Cells were lysed in RIPA buffer and quantified using the BCA method. Primary antibodies against GAPDH (2118, Cell Signaling Technology, 1:5,000), p70 S6 Kinase (2118, Cell Signaling Technology, 1:2,000), p-p70 S6 Kinase T389 (9234, Cell Signaling Technology, 1:1,000) were applied. After incubation with HRP-conjugated secondary antibody, membranes were developed with enhanced chemiluminescence western blotting substrate (32106, Thermo Fisher Scientific). Images were taken using an Amersham ImageQuant 800 biomolecular imager.

### Puromycin incorporation assay

To detect new protein synthesis, we performed the puromycin incorporation assay. For the *in vivo* assay, mice were treated with an intraperitoneal injection of puromycin (200µL per mouse, 2.5mM, HY-B1743, Med Chem Express) 1 hour before tissue collection. Newly synthesized proteins were detected by immunofluorescence staining using anti-puromycin antibody. For the *in vitro* assay, cells were treated with 20 µM O-propargyl-puromycin (OPP) for 30 minutes. The incorporated OPP was then detected using the Click-iT Plus OPP Alexa Fluor 647 Protein Synthesis Assay Kit (C10458, Thermo Fisher Scientific). Signals were detected using a BD LSRFortessa instrument.

### Data availability

Raw sequencing data of scRNA-seq is deposited to the Gene Expression Omnibus (GEO): GSE292432. All other data supporting the finding of this study are available upon request from the corresponding authors.

### Statistics

Statistical analysis was performed as detailed in the figure legends. All results were successfully repeated with a minimum of two independent experiments. Plots were generated using GraphPad Prism 10.

---

**Figure S1.**
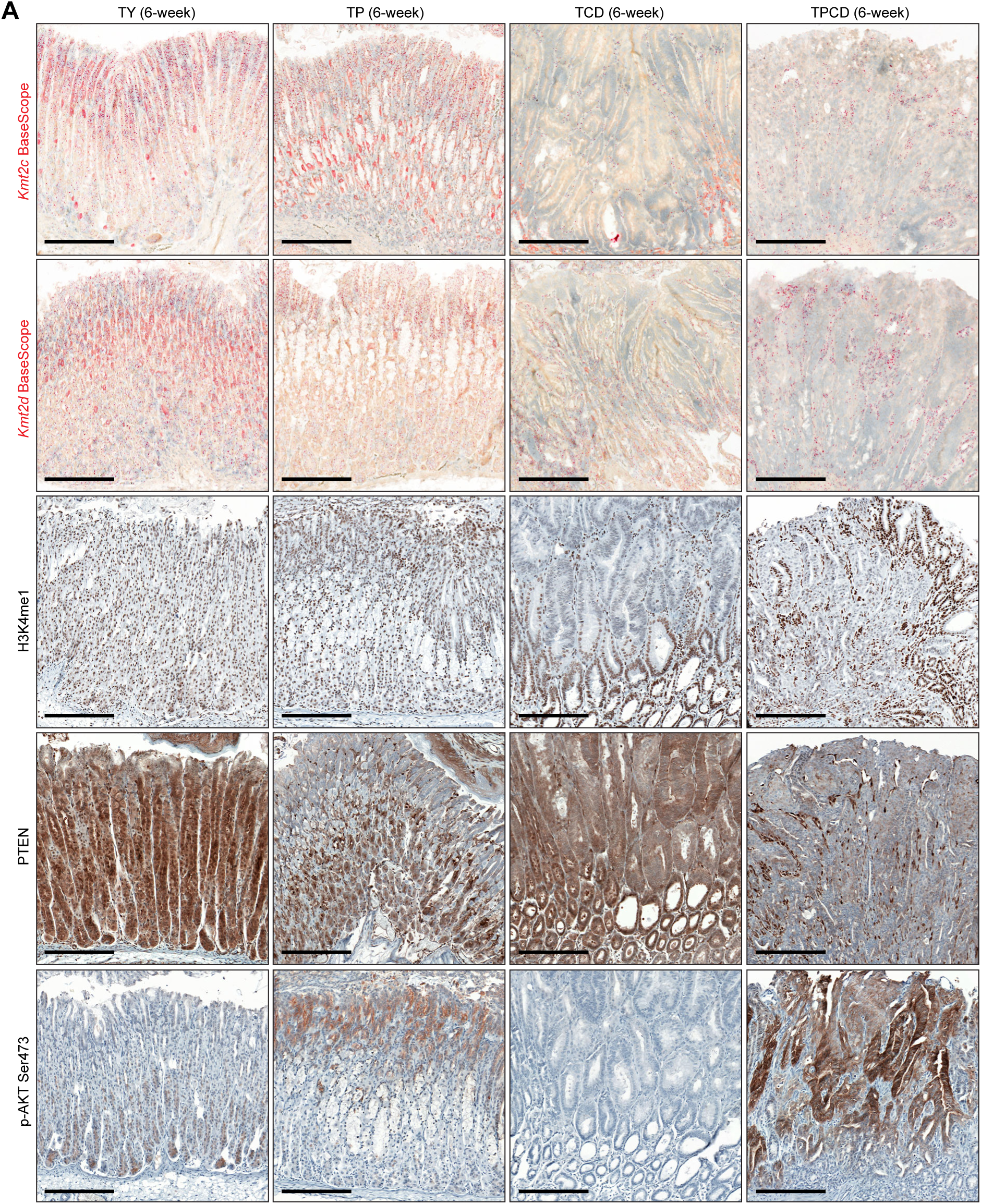
Validation of *Tmprss2-CreER^T2^*-mediated gene knockout in GEMMs. A, Top, representative BaseScope staining of stomach tissues with probes targeting *Kmt2c* or *Kmt2d* floxed exons. Bottom, representative IHC of H3K4me1, PTEN, and p-AKT Ser473 in stomach tissues. Scale bar, 200 µm.

**Figure S2.**
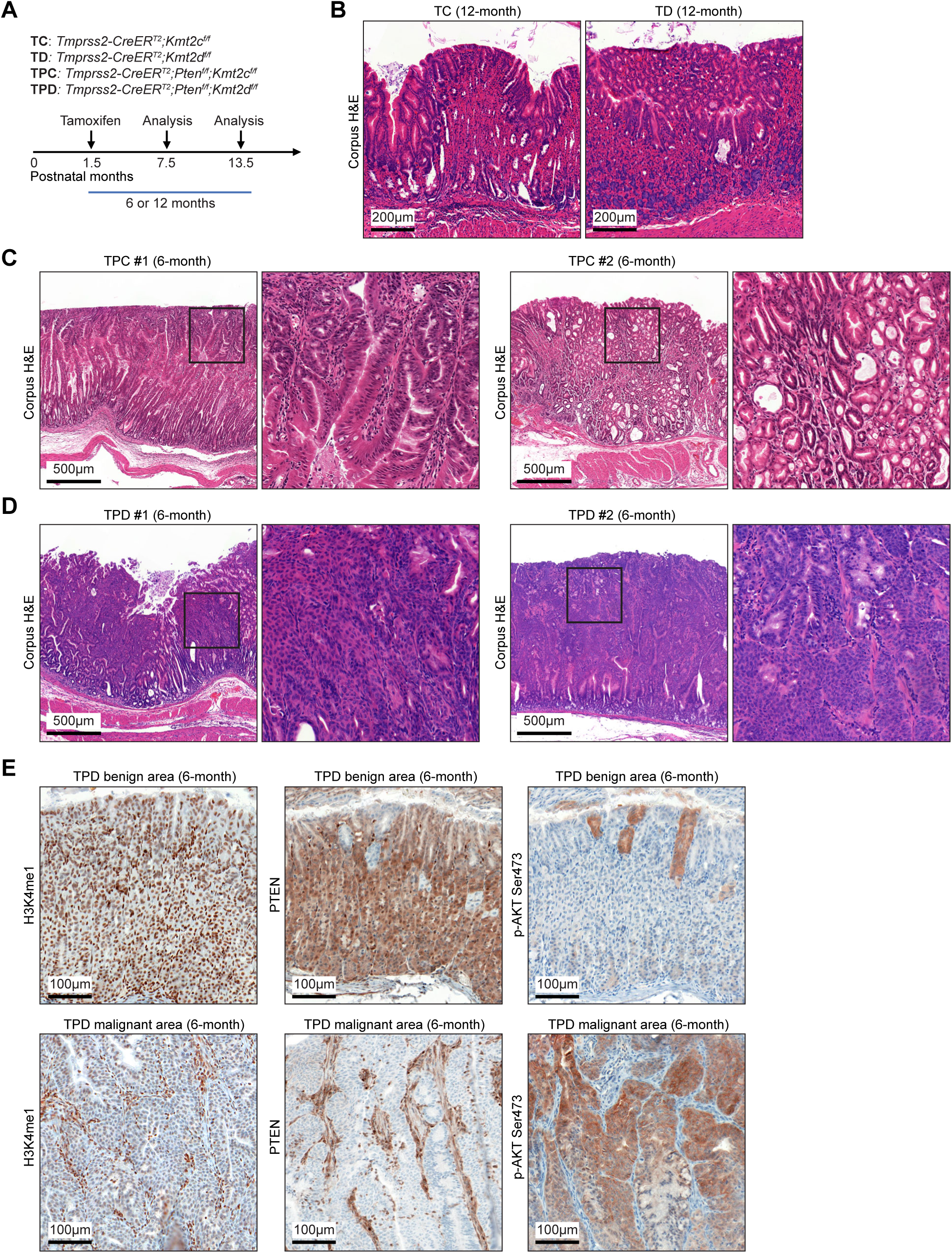
Deletion of both *Kmt2c* and *Kmt2d* is required for robust tumorigenesis. A, Schematic of mouse models: two doses of tamoxifen (3 mg × 2) were injected intraperitoneally with a 48-hour interval. B, Representative H&E staining of stomach tissues in TC and TD groups. Scale bar, 200 µm. C-D, Representative H&E staining of stomach tissues in TPC and TPD groups. Scale bar, 500 µm. E, Representative IHC of H3K4me1, PTEN, and p-AKT Ser473 in TPD stomach tissues. Scale bar, 100 µm.

**Figure S3.**
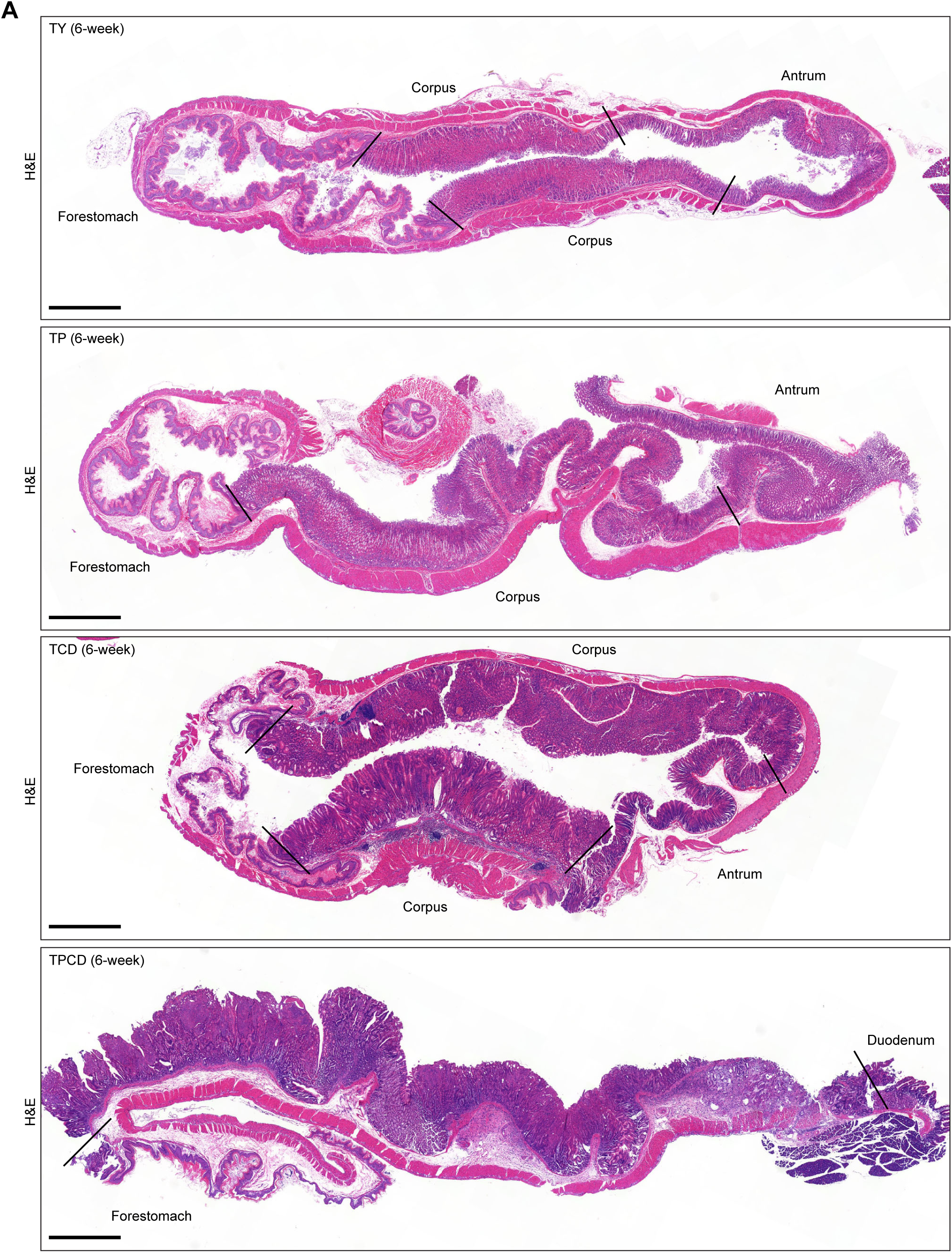
H&E staining in mouse stomach tissues. A, Representative H&E staining of stomach tissues after tamoxifen administration. Scale bar, 1 mm.

**Figure S4.**
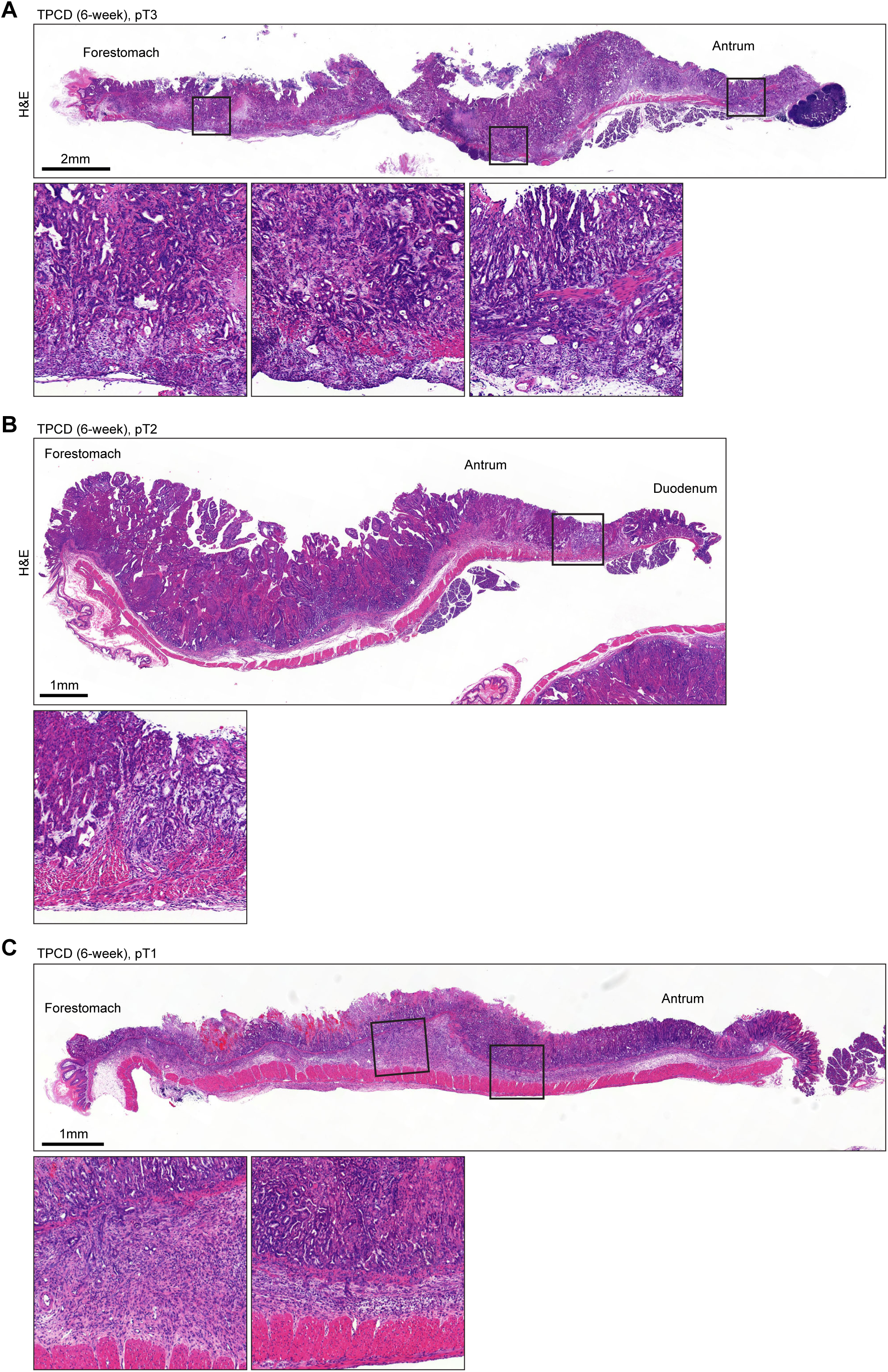
H&E staining showing muscle-invasive lesions in mouse stomach tissues. A-C, Representative H&E staining of pT1, pT2, and pT3 stages in TPCD stomach tissues.

**Figure S5.**
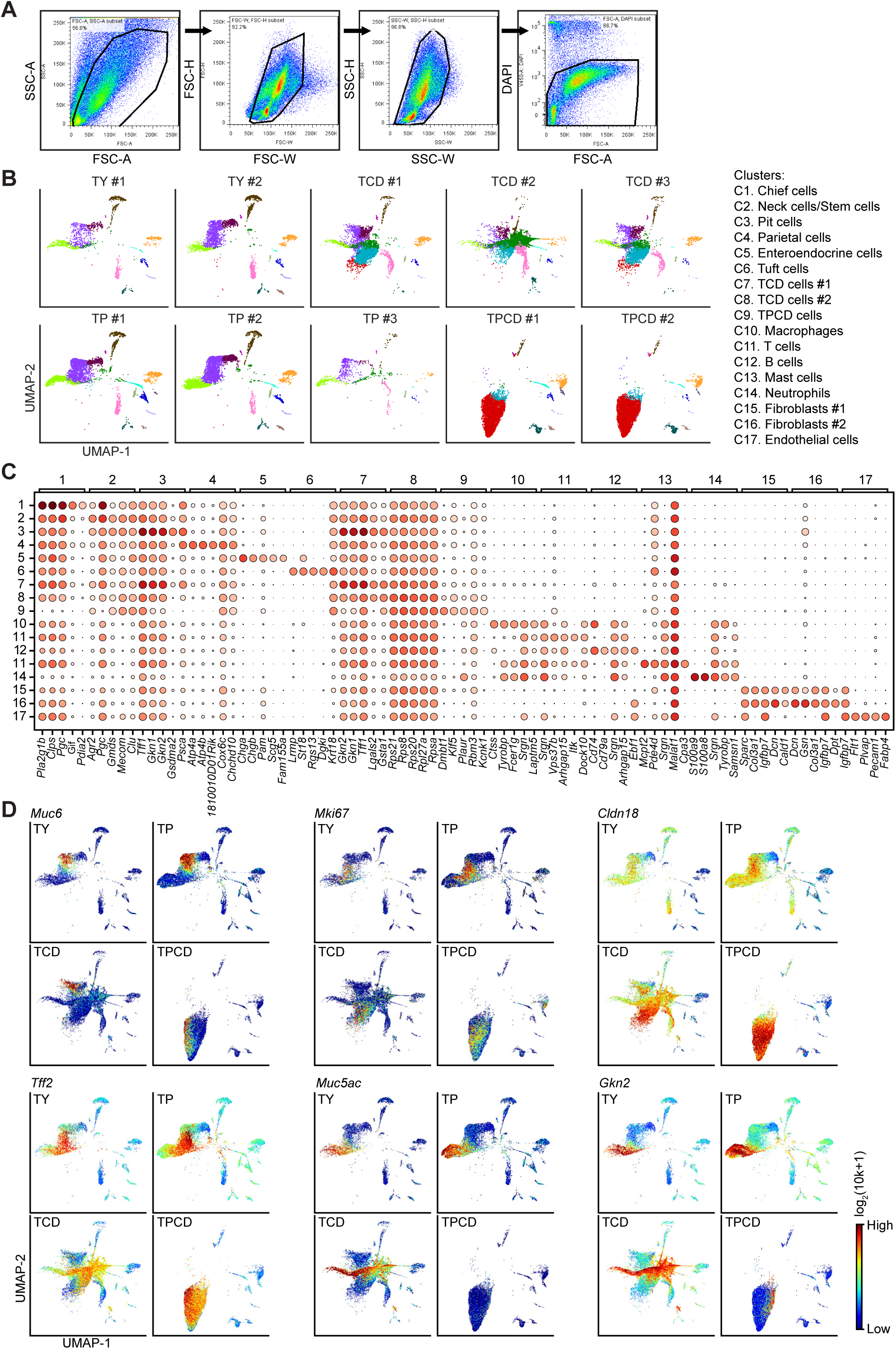
Characterization of cell clusters in scRNA-seq. A, Fluorescence-activated cell sorting (FACS) of DAPI-negative viable cells in dissociated stomach mucosa. B, UMAP showing reproducibility of clusters in TY, TP, TCD, and TPCD mice. C, Dot plot showing the expression of top 5 genes in each cluster. D, UMAP color-coded by expression of representative gastric lineage markers.

**Figure S6.**
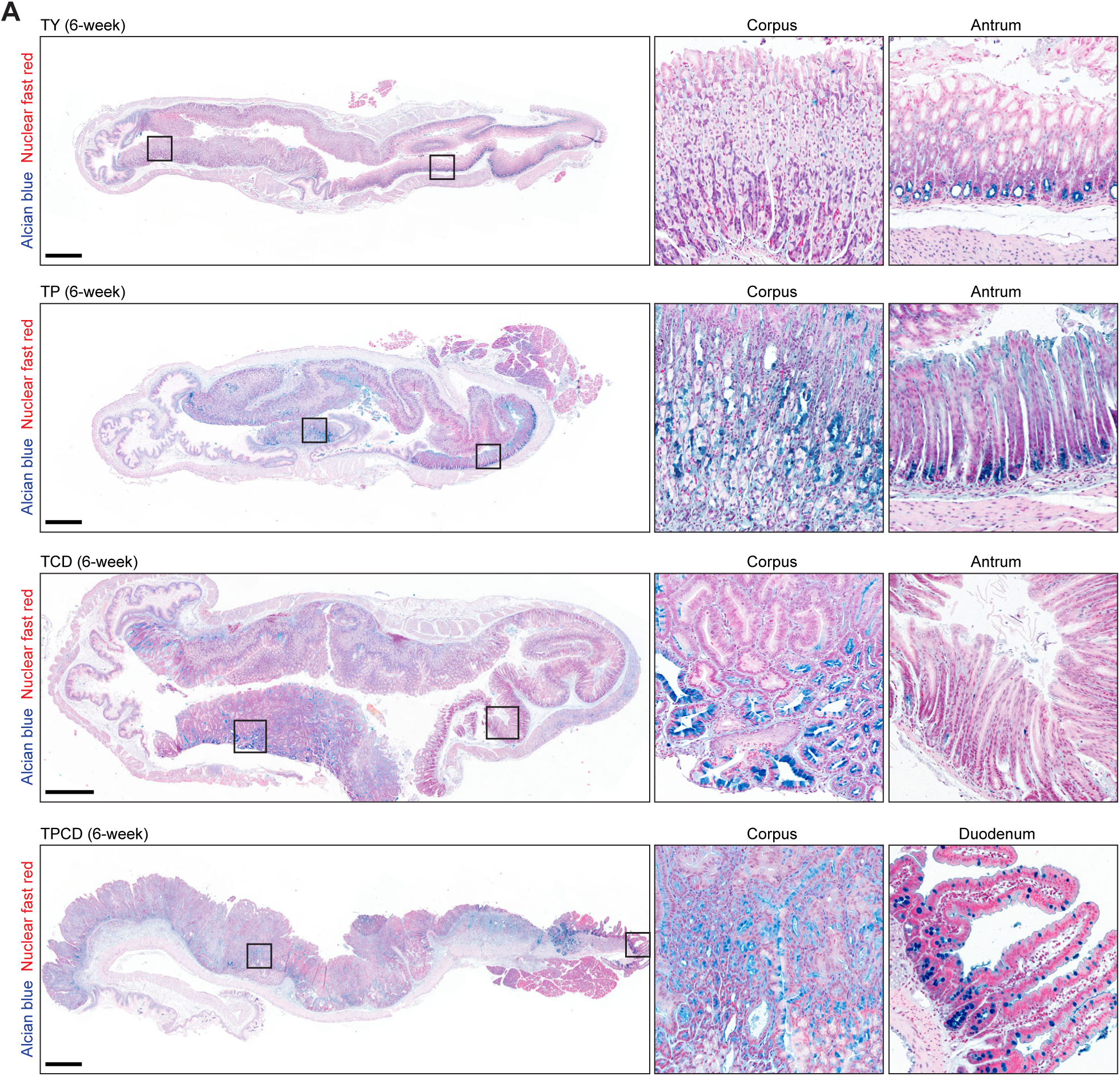
Alcian blue staining in mouse stomach tissues. A, Representative alcian blue staining in TY, TP, TCD, and TPCD stomach tissues. Nuclei were counterstained using nuclear fast red. Scale bar, 1 mm.

**Figure S7.**
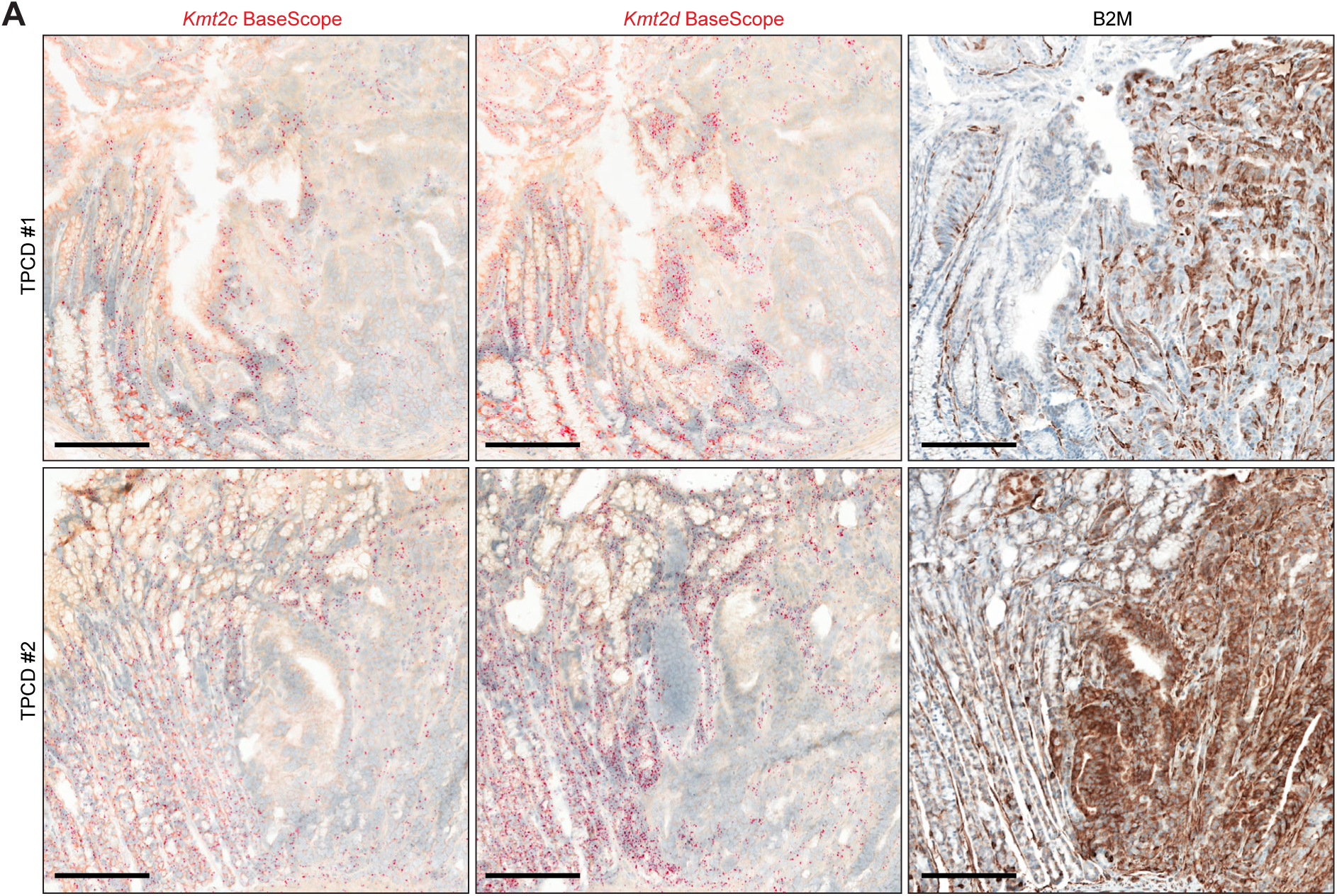
*Kmt2c/d* loss increases B2M protein levels. A, Representative BaseScope images of *Kmt2c* and *Kmt2d*, and representative IHC of B2M in TPCD stomach tissues. Note that *Kmt2c/d*-loss areas were negative for red dots, while *Kmt2c/d*-intact areas were positive for red dots. Scale bar, 200 µm.

**Figure S8.**
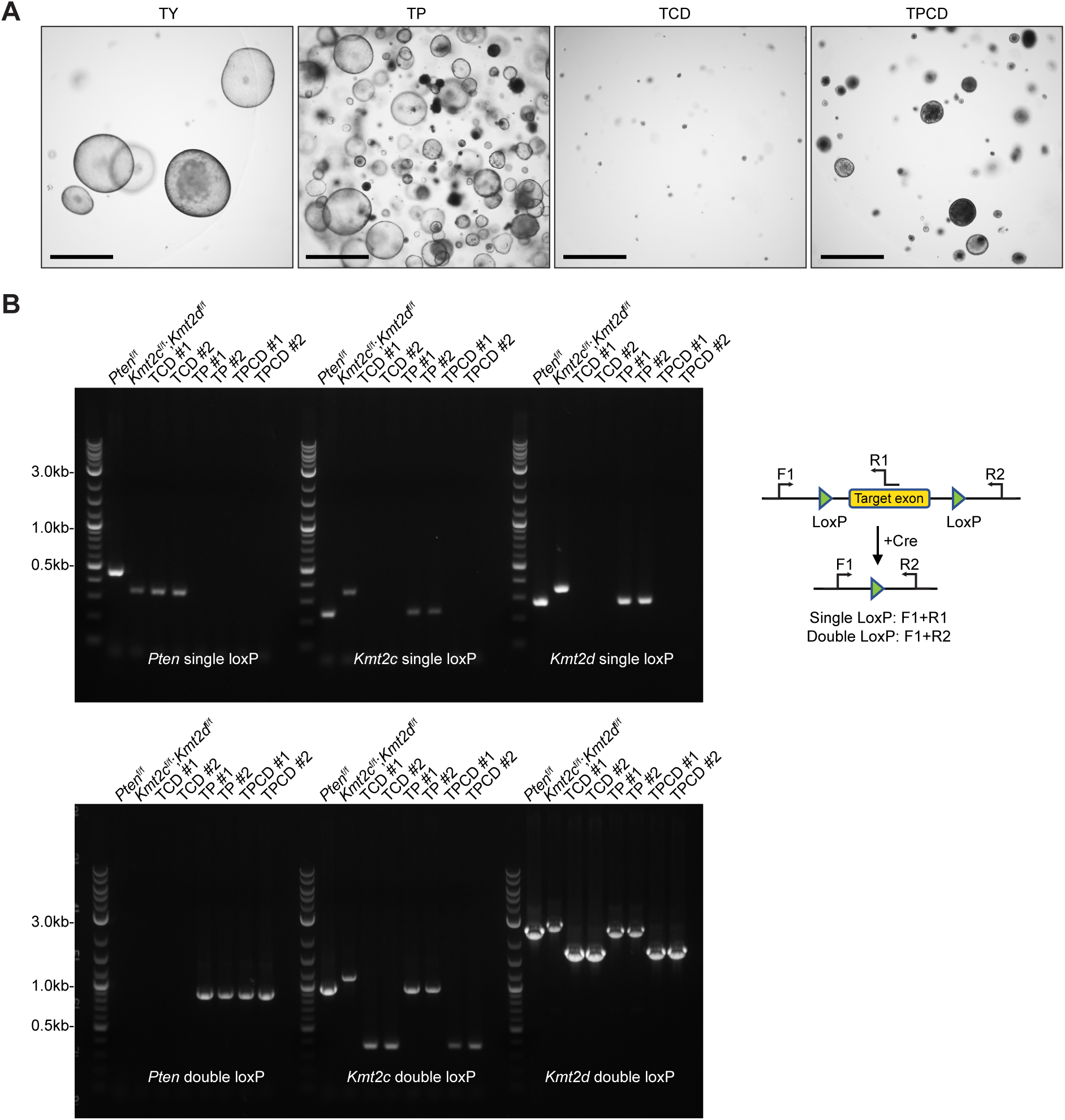
Validation of gene deletions in stomach organoids. A, Representative bright-field images of organoids in TY, TP, TCD, and TPCD groups. Scale bar, 1 mm. B, Genotyping of *Pten*, *Kmt2c*, and *Kmt2d* floxed alleles before and after 4-hydroxytamoxifen (4OHT) treatment.

**Figure S9.**
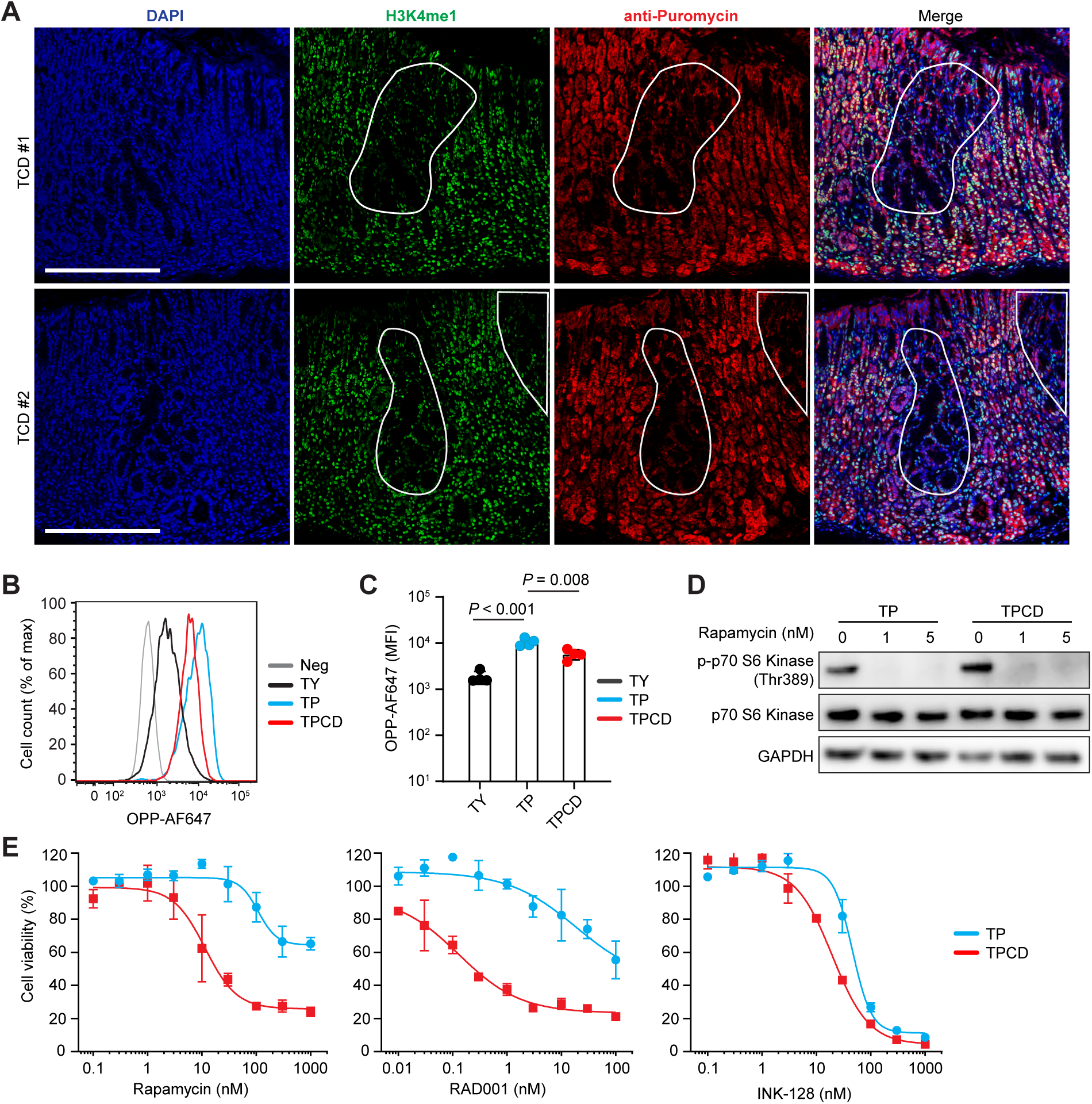
*Kmt2c/d* loss reduces new protein synthesis. A, Representative IF staining of H3K4me1 and Puromycin in TCD mice treated with puromycin. Cells in the highlighted area exhibited weaker staining of H3K4me1 and Puromycin. Nuclei were counterstained with DAPI. B-C, Flow cytometry analysis of OPP incorporation in TY, TP, and TPCD stomach cells. Data are presented as mean ± SD. Statistical analyses were performed with two-tailed t-test on log_10_ normalized data. D, Western blot analysis showing decreased phosphorylation of p70 S6 kinase (Thr389) with rapamycin treatment. TP and TPCD cells were treated for 1 h before sample collection. E, Cell viability following treatment of TP and TPCD cells with rapamycin, RAD-001, or INK-128 for 5 days.

**Figure S10.**
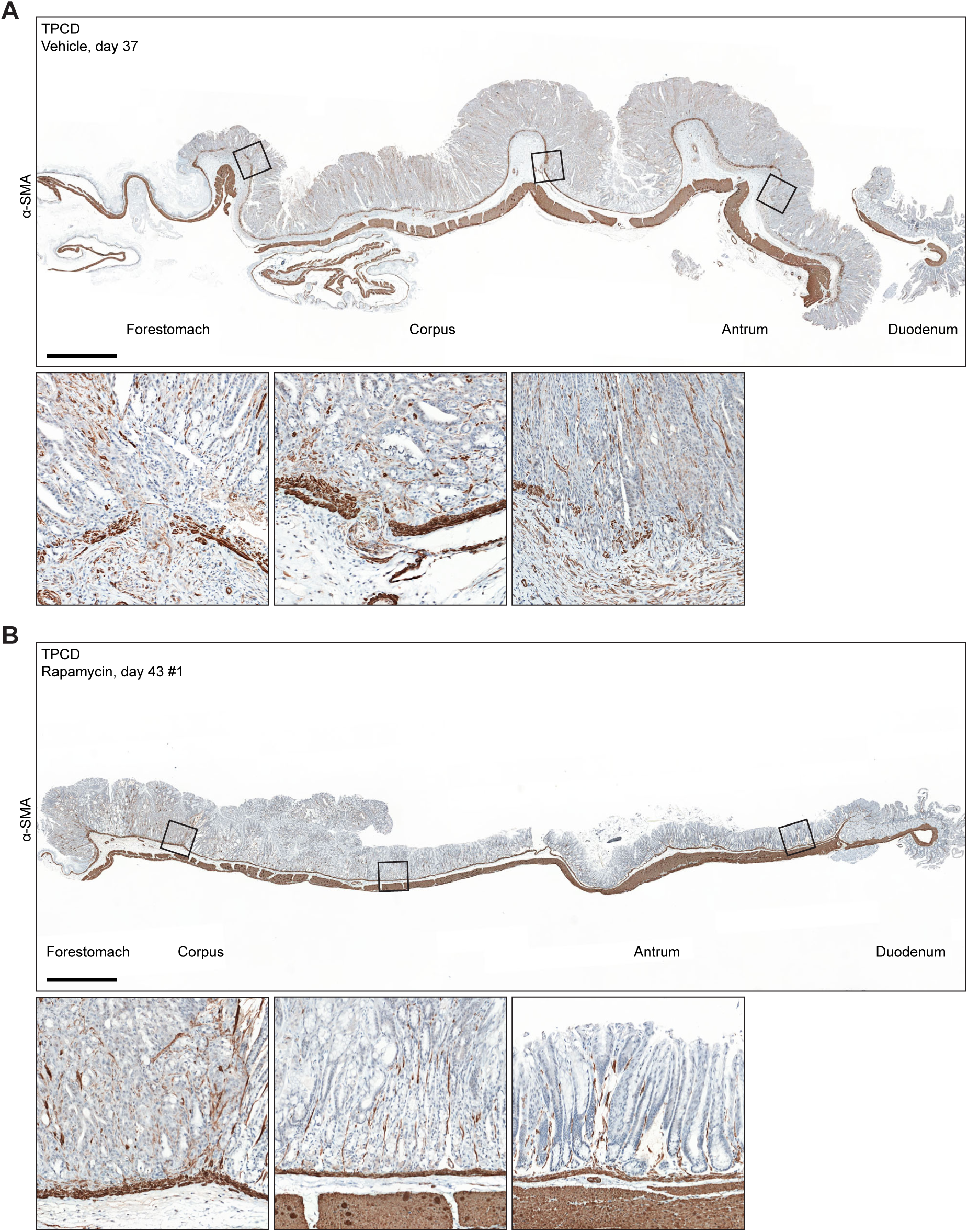
Rapamycin treatment reduces muscle-invasive lesions in TPCD mice. A, Representative IHC of α-SMA in TPCD stomach tissues treated with vehicle or rapamycin. Scale bar, 1 mm.

**Figure S11.**
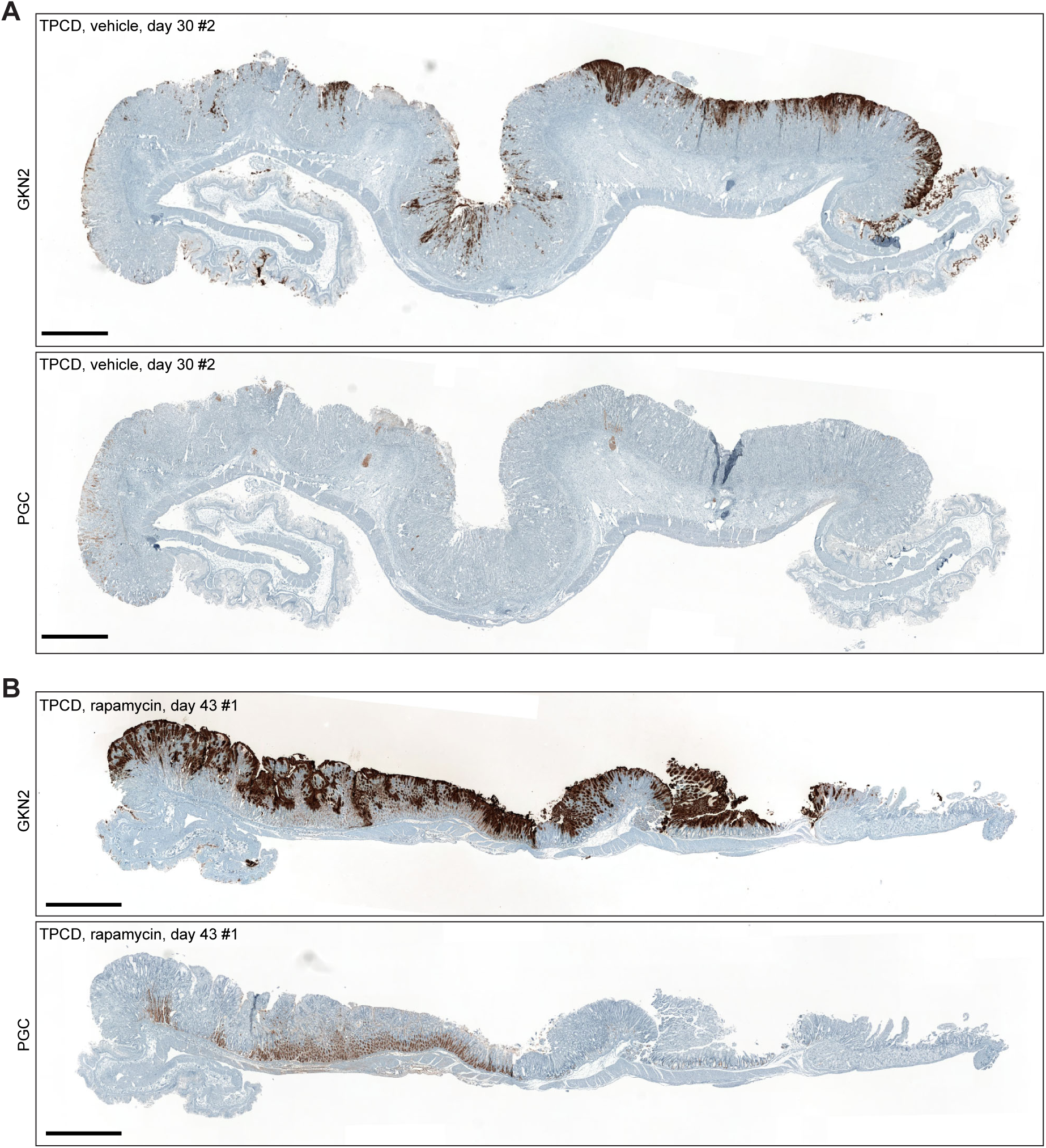
Rapamycin treatment rescues gastric differentiation in TPCD mice. A-B, Representative IHC of GKN2 and PGC in TPCD stomach tissues treated with vehicle or rapamycin. Scale bar, 1 mm.

## Reference

1 Sung H, Ferlay J, Siegel RL, Laversanne M, Soerjomataram I, Jemal A, Bray F. Global Cancer Statistics 2020: GLOBOCAN Estimates of Incidence and Mortality Worldwide for 36 Cancers in 185 Countries. CA Cancer J Clin 2021;71:209–49.

2 Smyth EC, Nilsson M, Grabsch HI, van Grieken NC, Lordick F. Gastric cancer. Lancet 2020;396:635–48.

3. 3 Cancer Genome Atlas Research N. Comprehensive molecular characterization of gastric adenocarcinoma. Nature 2014;513:202–9.

4 Cristescu R, Lee J, Nebozhyn M, Kim KM, Ting JC, Wong SS, et al. Molecular analysis of gastric cancer identifies subtypes associated with distinct clinical outcomes. Nat Med 2015;21:449–56.

5 Lauren P. The Two Histological Main Types of Gastric Carcinoma: Diffuse and So-Called Intestinal-Type Carcinoma. An Attempt at a Histo-Clinical Classification. Acta Pathol Microbiol Scand 1965;64:31–49.

6 Lo YH, Kolahi KS, Du Y, Chang CY, Krokhotin A, Nair A, et al. A CRISPR/Cas9-Engineered ARID1A-Deficient Human Gastric Cancer Organoid Model Reveals Essential and Nonessential Modes of Oncogenic Transformation. Cancer Discov 2021;11:1562–81.

7 Dong X, Song S, Li Y, Fan Y, Wang L, Wang R, et al. Loss of ARID1A activates mTOR signaling and SOX9 in gastric adenocarcinoma-rationale for targeting ARID1A deficiency. Gut 2022;71:467–78.

8 Piunti A, Shilatifard A. Epigenetic balance of gene expression by Polycomb and COMPASS families. Science 2016;352:aad9780.

9 Cenik BK, Shilatifard A. COMPASS and SWI/SNF complexes in development and disease. Nat Rev Genet 2021;22:38–58.

10 Wang N, Pachai MR, Li D, Lee CJ, Warda S, Khudoynazarova MN, et al. Loss of Kmt2c or Kmt2d primes urothelium for tumorigenesis and redistributes KMT2A-menin to bivalent promoters. Nat Genet 2025;57:165–79.

11 Hu D, Gao X, Morgan MA, Herz HM, Smith ER, Shilatifard A. The MLL3/MLL4 branches of the COMPASS family function as major histone H3K4 monomethylases at enhancers. Mol Cell Biol 2013;33:4745–54.

12 Alam H, Tang M, Maitituoheti M, Dhar SS, Kumar M, Han CY, et al. KMT2D Deficiency Impairs Super-Enhancers to Confer a Glycolytic Vulnerability in Lung Cancer. Cancer Cell 2020;37:599–617 e7.

13 Na F, Pan X, Chen J, Chen X, Wang M, Chi P, et al. KMT2C deficiency promotes small cell lung cancer metastasis through DNMT3A-mediated epigenetic reprogramming. Nat Cancer 2022;3:753–67.

14 Pan Y, Han H, Hu H, Wang H, Song Y, Hao Y, et al. KMT2D deficiency drives lung squamous cell carcinoma and hypersensitivity to RTK-RAS inhibition. Cancer Cell 2023;41:88–105 e8.

15 Seehawer M, Li Z, Nishida J, Foidart P, Reiter AH, Rojas-Jimenez E, et al. Loss of Kmt2c or Kmt2d drives brain metastasis via KDM6A-dependent upregulation of MMP3. Nat Cell Biol 2024;26:1165–75.

16 Ortega-Molina A, Boss IW, Canela A, Pan H, Jiang Y, Zhao C, et al. The histone lysine methyltransferase KMT2D sustains a gene expression program that represses B cell lymphoma development. Nat Med 2015;21:1199–208.

17 Wang J, Xiu J, Baca Y, Battaglin F, Arai H, Kawanishi N, et al. Large-scale analysis of KMT2 mutations defines a distinctive molecular subset with treatment implication in gastric cancer. Oncogene 2021;40:4894–905.

18 Zheng YC, Duan YC, Ma JL, Xu RM, Zi X, Lv WL, et al. Triazole-dithiocarbamate based selective lysine specific demethylase 1 (LSD1) inactivators inhibit gastric cancer cell growth, invasion, and migration. J Med Chem 2013;56:8543–60.

19 Wang DX, Long JY, Li RZ, Zhang DL, Liu H, Liu J, et al. Mutation status of the KMT2 family associated with immune checkpoint inhibitors (ICIs) therapy and implicating diverse tumor microenvironments. Mol Cancer 2024;23:15.

20. Cancer Genome Atlas N. Comprehensive molecular characterization of human colon and rectal cancer. Nature 2012;487:330–7.

21. Cancer Genome Atlas Research N, Kandoth C, Schultz N, Cherniack AD, Akbani R, Liu Y, et al. Integrated genomic characterization of endometrial carcinoma. Nature 2013;497:67–73.

22 Guo SL, Ye H, Teng Y, Wang YL, Yang G, Li XB, et al. Akt-p53-miR-365-cyclin D1/cdc25A axis contributes to gastric tumorigenesis induced by PTEN deficiency. Nat Commun 2013;4:2544.

23 Leibold J, Tsanov KM, Amor C, Ho YJ, Sanchez-Rivera FJ, Feucht J, et al. Somatic mouse models of gastric cancer reveal genotype-specific features of metastatic disease. Nat Cancer 2024;5:315–29.

24 Gao D, Zhan Y, Di W, Moore AR, Sher JJ, Guan Y, et al. A Tmprss2-CreERT2 Knock-In Mouse Model for Cancer Genetic Studies on Prostate and Colon. PLoS One 2016;11:e0161084.

25 Guo W, Li L, He J, Liu Z, Han M, Li F, et al. Single-cell transcriptomics identifies a distinct luminal progenitor cell type in distal prostate invagination tips. Nat Genet 2020;52:908–18.

26 Seidlitz T, Chen YT, Uhlemann H, Scholch S, Kochall S, Merker SR, et al. Mouse Models of Human Gastric Cancer Subtypes With Stomach-Specific CreERT2-Mediated Pathway Alterations. Gastroenterology 2019;157:1599–614 e2.

27 Huebner AJ, Gorelov RA, Deviatiiarov R, Demharter S, Kull T, Walsh RM, et al. Dissection of gastric homeostasis in vivo facilitates permanent capture of isthmus-like stem cells in vitro. Nat Cell Biol 2023;25:390–403.

28 Petersen CP, Mills JC, Goldenring JR. Murine Models of Gastric Corpus Preneoplasia. Cell Mol Gastroenterol Hepatol 2017;3:11–26.

29 Franzen O, Gan LM, Bjorkegren JLM. PanglaoDB: a web server for exploration of mouse and human single-cell RNA sequencing data. Database (Oxford) 2019;2019.

30 Busslinger GA, Weusten BLA, Bogte A, Begthel H, Brosens LAA, Clevers H. Human gastrointestinal epithelia of the esophagus, stomach, and duodenum resolved at single-cell resolution. Cell Rep 2021;34:108819.

31 Quante M, Marrache F, Goldenring JR, Wang TC. TFF2 mRNA transcript expression marks a gland progenitor cell of the gastric oxyntic mucosa. Gastroenterology 2010;139:2018–27 e2.

32 Nakayama I, Qi C, Chen Y, Nakamura Y, Shen L, Shitara K. Claudin 18.2 as a novel therapeutic target. Nat Rev Clin Oncol 2024;21:354–69.

33 Correa P, Piazuelo MB, Wilson KT. Pathology of gastric intestinal metaplasia: clinical implications. Am J Gastroenterol 2010;105:493–8.

34 Barros R, Freund JN, David L, Almeida R. Gastric intestinal metaplasia revisited: function and regulation of CDX2. Trends Mol Med 2012;18:555–63.

35 Zhang P, Yang M, Zhang Y, Xiao S, Lai X, Tan A, et al. Dissecting the Single-Cell Transcriptome Network Underlying Gastric Premalignant Lesions and Early Gastric Cancer. Cell Rep 2019;27:1934–47 e5.

36 Kumar V, Ramnarayanan K, Sundar R, Padmanabhan N, Srivastava S, Koiwa M, et al. Single-Cell Atlas of Lineage States, Tumor Microenvironment, and Subtype-Specific Expression Programs in Gastric Cancer. Cancer Discov 2022;12:670–91.

37 Korsunsky I, Millard N, Fan J, Slowikowski K, Zhang F, Wei K, et al. Fast, sensitive and accurate integration of single-cell data with Harmony. Nat Methods 2019;16:1289–96.

38 Wang G, Chow RD, Zhu L, Bai Z, Ye L, Zhang F, et al. CRISPR-GEMM Pooled Mutagenic Screening Identifies KMT2D as a Major Modulator of Immune Checkpoint Blockade. Cancer Discov 2020;10:1912–33.

39 Hsieh AC, Liu Y, Edlind MP, Ingolia NT, Janes MR, Sher A, et al. The translational landscape of mTOR signalling steers cancer initiation and metastasis. Nature 2012;485:55–61.

40 Ando S, Perkins CM, Sajiki Y, Chastain C, Valanparambil RM, Wieland A, et al. mTOR regulates T cell exhaustion and PD-1-targeted immunotherapy response during chronic viral infection. J Clin Invest 2023;133.

41 Lu Z, Zhong A, Liu H, Zhang M, Chen X, Pan X, et al. Dissecting the genetic and microenvironmental factors of gastric tumorigenesis in mice. Cell Rep 2022;41:111482.

42 Je EM, Lee SH, Yoo NJ, Lee SH. Mutational and expressional analysis of MLL genes in gastric and colorectal cancers with microsatellite instability. Neoplasma 2013;60:188–95.

43 Cortes-Ciriano I, Lee S, Park WY, Kim TM, Park PJ. A molecular portrait of microsatellite instability across multiple cancers. Nat Commun 2017;8:15180.

44 Nowicki-Osuch K, Zhuang L, Cheung TS, Black EL, Masque-Soler N, Devonshire G, et al. Single-Cell RNA Sequencing Unifies Developmental Programs of Esophageal and Gastric Intestinal Metaplasia. Cancer Discov 2023;13:1346–63.

45 Huang KK, Ma H, Chong RHH, Uchihara T, Lian BSX, Zhu F, et al. Spatiotemporal genomic profiling of intestinal metaplasia reveals clonal dynamics of gastric cancer progression. Cancer Cell 2023;41:2019–37 e8.

46 Kumagai K, Shimizu T, Takai A, Kakiuchi N, Takeuchi Y, Hirano T, et al. Expansion of Gastric Intestinal Metaplasia with Copy Number Aberrations Contributes to Field Cancerization. Cancer Res 2022;82:1712–23.

47. Shitara K, Lordick F, Bang YJ, Enzinger P, Ilson D, Shah MA, et al. Zolbetuximab plus mFOLFOX6 in patients with CLDN18.2-positive, HER2-negative, untreated, locally advanced unresectable or metastatic gastric or gastro-oesophageal junction adenocarcinoma (SPOTLIGHT): a multicentre, randomised, double-blind, phase 3 trial. Lancet 2023;401:1655–68.

48 Shah MA, Shitara K, Ajani JA, Bang YJ, Enzinger P, Ilson D, et al. Zolbetuximab plus CAPOX in CLDN18.2-positive gastric or gastroesophageal junction adenocarcinoma: the randomized, phase 3 GLOW trial. Nat Med 2023;29:2133–41.

49 Liu L, Niu L, Zheng X, Xiao F, Sun H, Deng W, Cai J. PD-L1 expression-related PI3K pathway correlates with immunotherapy efficacy in gastric cancer. Ther Adv Med Oncol 2023;15:17588359231205853.

50 Takei S, Kawazoe A, Shitara K. The New Era of Immunotherapy in Gastric Cancer. Cancers (Basel) 2022;14.

51 Ohtsu A, Ajani JA, Bai YX, Bang YJ, Chung HC, Pan HM, et al. Everolimus for previously treated advanced gastric cancer: results of the randomized, double-blind, phase III GRANITE-1 study. J Clin Oncol 2013;31:3935–43.

52 Moore EC, Cash HA, Caruso AM, Uppaluri R, Hodge JW, Van Waes C, Allen CT. Enhanced Tumor Control with Combination mTOR and PD-L1 Inhibition in Syngeneic Oral Cavity Cancers. Cancer Immunol Res 2016;4:611–20.

53 Mathew D, Marmarelis ME, Foley C, Bauml JM, Ye D, Ghinnagow R, et al. Combined JAK inhibition and PD-1 immunotherapy for non-small cell lung cancer patients. Science 2024;384:eadf1329.

54 Li D, Zhan Y, Wang N, Tang F, Lee CJ, Bayshtok G, et al. ETV4 mediates dosage-dependent prostate tumor initiation and cooperates with p53 loss to generate prostate cancer. Sci Adv 2023;9:eadc9446.

55 Wolf FA, Angerer P, Theis FJ. SCANPY: large-scale single-cell gene expression data analysis. Genome Biol 2018;19:15.

56 Traag VA, Waltman L, van Eck NJ. From Louvain to Leiden: guaranteeing well-connected communities. Sci Rep 2019;9:5233.

57 Baumgartner CK, Ebrahimi-Nik H, Iracheta-Vellve A, Hamel KM, Olander KE, Davis TGR, et al. The PTPN2/PTPN1 inhibitor ABBV-CLS-484 unleashes potent anti-tumour immunity. Nature 2023;622:850–62.

